# Predictive engineering and optimization of tryptophan metabolism in yeast through a combination of mechanistic and machine learning models

**DOI:** 10.1101/858464

**Authors:** Jie Zhang, Søren D. Petersen, Tijana Radivojevic, Andrés Ramirez, Andrés Pérez, Eduardo Abeliuk, Benjamín J. Sánchez, Zachary Costello, Yu Chen, Mike Fero, Hector Garcia Martin, Jens Nielsen, Jay D. Keasling, Michael K. Jensen

## Abstract

In combination with advanced mechanistic modeling and the generation of high-quality multi-dimensional data sets, machine learning is becoming an integral part of understanding and engineering living systems. Here we show that mechanistic and machine learning models can complement each other and be used in a combined approach to enable accurate genotype-to-phenotype predictions. We use a genome-scale model to pinpoint engineering targets and produce a large combinatorial library of metabolic pathway designs with different promoters which, once phenotyped, provide the basis for machine learning algorithms to be trained and used for new design recommendations. The approach enables successful forward engineering of aromatic amino acid metabolism in yeast, with the new recommended designs improving tryptophan production by up to 17% compared to the best designs used for algorithm training, and ultimately producing a total increase of 106% in tryptophan accumulation compared to optimized reference designs. Based on a single high-throughput data-generation iteration, this study highlights the power of combining mechanistic and machine learning models to enhance their predictive power and effectively direct metabolic engineering efforts.

## INTRODUCTION

Metabolic engineering is the directed improvement of cell properties through the modification of specific biochemical reactions (Stephanopoulos, 1999). Beyond offering an improved understanding of basic cellular metabolism, the field of metabolic engineering also envisions sustainable production of biomolecules for health, food, and manufacturing industries, by fermenting feedstocks into value-added biomolecules using engineered cells (Keasling, 2010). These promises leverage tools and technologies developed over recent decades which include mechanistic metabolic modeling, targeted genome engineering, and robust bioprocess optimization; ultimately aiming for accurate and scalable predictions of cellular phenotypes from deduced genotypes (Nielsen and Keasling, 2016; Choi et al., 2019; Liu and Nielsen, 2019).

Among the different types of mechanistic models for simulating metabolism, genome-scale models (GSMs) are one of the most popular approaches, as they are genome-complete, covering thousands of metabolic reactions. These computational models not only provide qualitative mapping of cellular metabolism (Hefzi et al., 2016; Monk et al., 2017; Lu et al., 2019), but have also been successfully applied for the discovery of novel metabolic functions (Guzmán et al., 2015), and to guide engineering designs towards desired phenotypes (Yang et al., 2018).As GSMs are built based only on the stoichiometry of metabolic reactions, several methods have been developed to account for additional layers of information regarding the chemical intermediates and the catalyzing enzymes participating in the metabolic pathways of interest (Lewis et al., 2012). However, the predictive power of these enhanced models is often hampered by the limited knowledge and data available for any of such parameters affecting metabolic regulation (Gardner, 2013; Khodayari et al., 2015; Long and Antoniewicz, 2019).

Machine learning provides a complementary approach to guide metabolic engineering by learning patterns on systems behavior from large experimental data sets (Camacho et al., 2018). As such, machine learning models differ from mechanistic models by being purely data-driven. Indeed, machine learning methods for the generation of predictive models on living systems are becoming ubiquitous, including applications within genome annotation, *de novo* pathway discovery, product maximization in engineered microbial cells, pathway dynamics, and transcriptional drivers of disease states (Alonso-Gutierrez et al., 2015; Carro et al., 2010; Costello and Martin, 2018; Jervis et al., 2019; Mellor et al., 2016; Schläpfer et al., 2017). While being able to provide predictive power based on complex multivariate relationships (Presnell and Alper, 2019), the training of machine learning algorithms requires large datasets of high quality, and thereby imposes certain standards for the experimental workflows. For instance, for genotype-to-phenotype predictions, it is desirable that datasets contain a high variation between both genotypes and phenotypes (Carbonell et al., 2019). Also, measurements on the individual experimental unit, e.g. a strain, should be accurate and obtainable in a high-throughput manner, in order to limit the number of iterative design-build-test cycles needed in order to reach the desired output.

While mechanistic models require *a priori* knowledge of the living system of interest, and machine learning-guided predictions require ample multivariate experimental data for training, the combination of mechanistic and machine learning models holds promise for improved performance of predictive engineering of cells by uniting the advantages of the causal understanding of mechanism from mechanistic models with the predictive power of machine learning (Zampieri et al., 2019; Presnell and Alper, 2019). Metabolic pathways are known to be regulated at multiple levels, including transcriptional, translational, and allosteric levels (Chubukov et al., 2014). To cost-effectively move through the design and build steps of complex metabolic pathways regulated at multiple levels, combinatorial optimization of metabolic pathways, in contrast to sequential genotype edits, has been demonstrated to effectively facilitate identification of global optima for outputs of interest (i.e. production; Jeschek et al., 2017). Searching global optima using combinatorial approaches involves facing an exponentially growing number of designs (known as the combinatorial explosion), and requires efficient building of multi-parameterized combinatorial libraries. However, this challenge can be mitigated by the use of intelligently designed condensed libraries which allow uniform discretisation of multidimensional spaces: e.g. by using well-characterized sets of DNA elements controlling the expression of candidate genes at defined levels (Jeschek et al., 2016; Lee et al., 2013). As cellular metabolism is regulated at multiple levels (Feng et al., 2014; Lahtvee et al., 2017), an efficient search strategy for global optima using combinatorial approaches should also take this into consideration, e.g. by using mechanistic models, ‘omics data repositories, and *a priori* biological understanding.

Here we combine mechanistic and machine learning models to enable robust genotype-to-phenotype predictions as a tool for metabolic engineering. The approach is exemplified for predictive engineering and optimization of the complexly regulated aromatic amino acid pathway that produces tryptophan in baker’s yeast *Saccharomyces cerevisiae*. We defined a 7,776-membered combinatorial library design space, based on 5 genes selected from GSM simulations and *a priori* biological understanding, each controlled at the level of gene expression by 6 different promoters from a total set of 30 promoters selected from transcriptomics data mining. In order to train predictive models for high-tryptophan biosynthesis rate in yeast, we collected >144,000 experimental data points using a tryptophan biosensor, exploring this way approximately 4% of the genetic designs of the library design space. Based on a single Design-Build-Test-Learn cycle focused on sequencing data, growth profiles, and biosensor output, we trained various machine learning algorithms. Predictive models based on these algorithms enabled construction of designs exhibiting tryptophan biosynthesis rates 106% higher than a state-of-the-art high-tryptophan reference strain (Hartmann et al., 2003; Rodriguez et al., 2015), and up to 17% higher rate than best designs used for training the models.

## RESULTS

### Model-guided design of high tryptophan production

One prime example of the multi-tiered complexity regulating metabolic fluxes, is the shikimate pathway, driving the central metabolic route leading to aromatic amino acid biosynthesis in microorganisms (Lingens et al., 1967; Braus, 1991; Averesch and Krömer, 2018). This pathway has enormous industrial relevance, since it has been used to produce bio-based replacements of a wealth of fossil fuel-derived aromatics, polymers, and potent human therapeutics (Curran et al., 2013; Suástegui and Shao, 2016).

To search for gene targets predicted to perturb tryptophan production, we initially performed constraint-based modeling for predicting single gene targets, with a simulated objective of combining growth and tryptophan production (Orth et al., 2010; Ferreira et al., 2019). From this analysis, we retrieved 192 genes, covering 259 biochemical reactions, that showed considerable changes as production shifted from growth towards tryptophan production (Figure 1A-B, Table S4). By performing an analysis for statistical over-representation of genome-scale modelled metabolic pathways, we observed that both the pentose phosphate pathway and glycolysis were among the top pathways with a significantly higher number of gene targets compared to the representation of all metabolic genes (Figure 1C, Table S5). Among the predicted gene targets in those pathways, *CDC19, TKL1, TAL1* and *PCK1* were initially selected as targets for combinatorial library construction (Figure 1B), as these genes have all been experimentally validated to be directly linked or to have an indirect impact on the shikimate pathway precursors erythrose 4-phosphate (E4P) and phosphoenolpyruvate (PEP). Specifically, *CDC19* encodes the major isoform of pyruvate kinase converting PEP into pyruvate to fuel the tricarboxylic acid (TCA) cycle, while *TKL1* and *TAL1* that encode the major isoform of transketolase and transaldolase, respectively, in the reversible non-oxidative pentose phosphate pathway (PPP), have been reported to impact the supply of E4P (Patnaik and Liao, 1994; Curran et al., 2013). Additionally, focusing on the E4P and PEP linkage, *PCK1* encoding PEP carboxykinase, was also selected due to its regeneration capacity of PEP from oxaloacetate (Yin, 1996). Lastly, while not being predicted as a target by the constraint-based modeling approach, the *PFK1* gene, encoding the alpha subunit of heterooctameric phosphofructokinase, catalyzing the irreversible conversion of fructose 6-phosphate (F6P) to fructose 1,6-bisphosphate (FBP), was selected, as insufficient activity of this enzyme is known to cause divergence of carbon flux towards the pentose phosphate pathway in different organisms across different kingdoms (Wang et al., 2013; Zhang et al., 2016).

**Figure 1.**
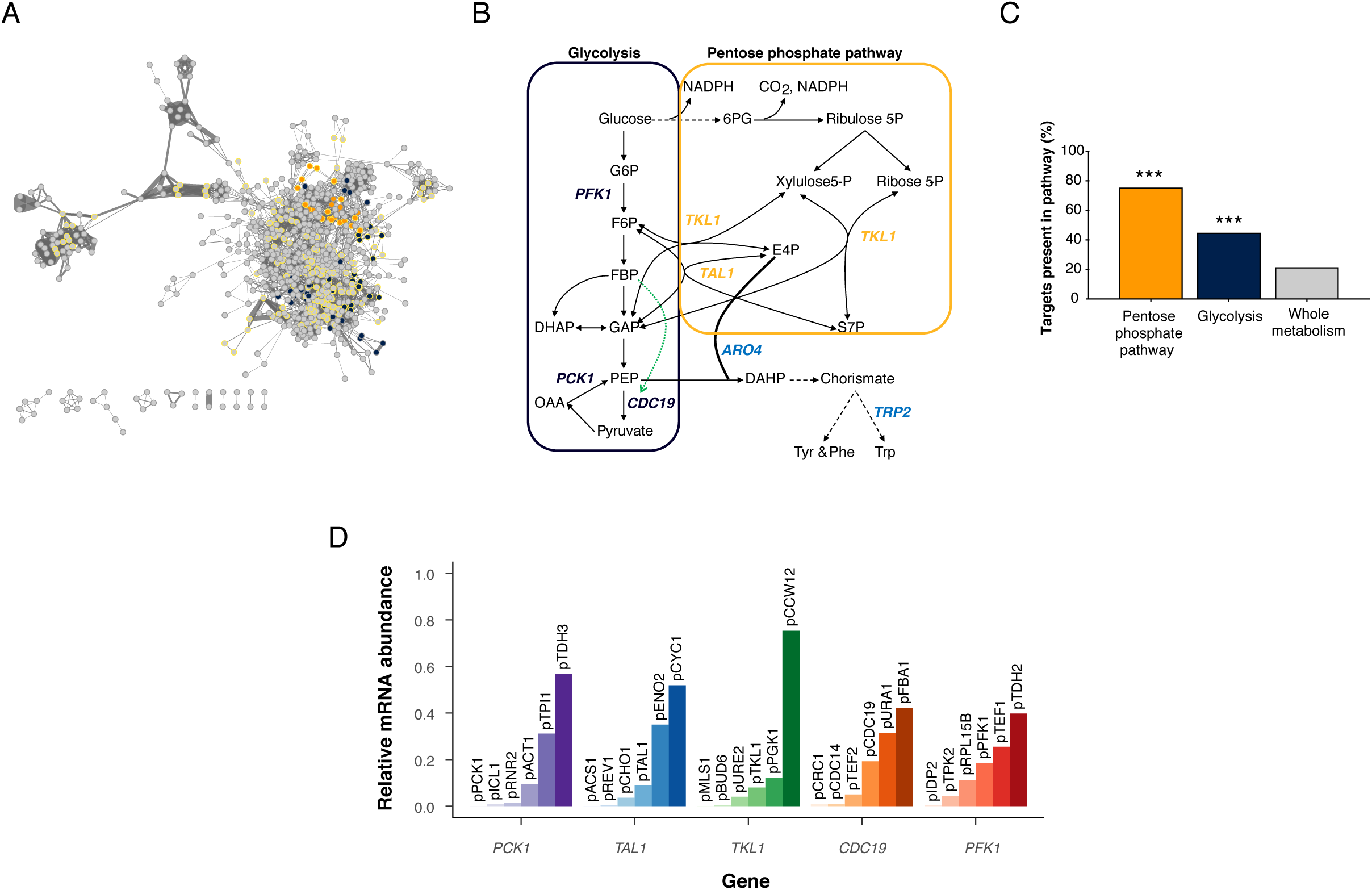
Selection of gene targets and promoters for combinatorial engineering of tryptophan metabolism in *S. cerevisiae*. (A) Gene-gene interaction network built with Cytoscape (Shannon et al., 2003), showing that pentose phosphate pathway and glycolysis are both in the core of metabolism in close proximity to many genes. Nodes are all 909 genes in yeast metabolism (Aung et al., 2013), sharing connections based on the number of shared metabolites by the corresponding reactions that the genes are related to: the thicker the edge, the higher the number of shared metabolites. Currency metabolites such as water, protons, ATP, etc. are removed from the analysis. The prefuse force directed layout is used for displaying the network. Genes are highlighted with a yellow border if they are selected targets by the mechanistic modeling approach, and in orange and dark blue if they belong to the pentose phosphate pathway or glycolysis, respectively. (B) Simplified map of metabolism showing the selected gene targets from glycolysis (dark blue) and pentose phosphate pathway (orange) based on a combination of mechanistic genome-scale modeling and literature studies for optimizing tryptophan production. Black dashed lines indicate multi-step reactions. Dashed green line indicates allosteric activation. G6P, glucose 6-phosphate; F6P, fructose 6-phosphate; FBP, fructose 1,6-bisphosphate; GAP, glyceraldehyde 3-phosphate; DHAP, dihydroxyacetone phosphate; PEP, phosphoenolpyruvate; OAA, oxaloacetate; 6PG, 6-phosphogluconate; E4P, erythrose 4-phosphate; S7P, sedoheptulose 7-phosphate; DAHP, 3-deoxy-7-phosphoheptulonate; Tyr, tyrosine; Phe, phenylalanine; Trp, tryptophan. (C) Percentage of genes in glycolysis (dark blue) and pentose phosphate pathway (orange) that were predicted by the mechanistic modelling to increase tryptophan production compared to the percentage of genes predicted as targets from the whole metabolism. *** = P-value < 0.05, Fisher’s exact testing. (D) Relative mRNA abundance, calculated for each gene as the proportion of mRNA reads obtained for any given promoter relative to the total sum of mRNA reads from each bin of six promoters. Absolute abundances for the 30 promoters were measured in S. cerevisiae CEN.PK 113-7D in the mid-log phase (Rajkumar et al., 2019). The promoters are grouped according to intended combinatorial gene associations.

Next, we mined transcriptomics data sets for the selection of promoters to control the expression of the five selected candidate genes. Here we focused on well-characterized and sequence-diverse promoters, to ensure rational designs spanning large absolute levels of promoter activities and limit the risk of recombination within strain designs and loss of any genetic elements, respectively (Figure S1; Rajkumar et al., 2019; Reider Apel et al., 2017). Together, this mining resulted in the selection of 25 sequence-diverse promoters, which together with the five promoters natively regulating the selected candidate genes, constitutes the parts catalog for combinatorial library design (Figure 1D; Figure S1, Table S6).

### Creation of a platform strain for a combinatorial library

To construct a combinatorial library targeting equal representation of thirty promoters expressing five candidate genes, we harnessed high-fidelity homologous recombination in yeast together with the targetability of CRISPR/Cas9 genome engineering for a one-pot assembly of a maximum of 7,776 (6^5^) different combinatorial designs. Due to the dramatic decrease in transformation efficiency when simultaneously targeting multiple loci in the genome (Jakočiu□nas et al., 2015), we targeted the sequential deletion of all five selected target genes from their original genomic loci, and next assemble a cluster of five expression cassettes into a single genomic landing as recently successfully reported for the “single-locus glycolysis” in yeast (Kuijpers et al., 2016)(Figure 2A). However, as *CDC19* is an essential gene, and deletion of *PFK1* causes growth retardation (Breslow et al., 2008; Cherry et al., 2012), this genetic background would be unsuitable for efficient one-pot transformation. For this reason our platform strain for library construction had a galactose-curable plasmid introduced expressing *PFK1, CDC19, TKL1* and *TAL1* under their native promoters (see METHODS DETAILS), before performing two sequential rounds of genome engineering to delete *PCK1, TKL1* and *TAL1*, and knock-down *CDC19* and *PFK1* using the weak promoters *RNR2* and *REV1*, respectively (Figure 2A). Furthermore, prior to one-pot assembly of the combinatorial library, we integrated the two feedback-inhibited shikimate pathway enzymes 3-deoxy-D-arabinose-heptulosonate-7-phosphate (DAHP) synthase (ARO4^K229L^) and anthranilate synthase (TRP2^S65R, S76L^) into our platform strain (Hartmann et al., 2003; Graf et al., 1993), thereby aiming to maximise the impact from transcriptional regulation of candidate genes on the overall tryptophan output, as removal of allosteric feedback inhibition is known to increase amino acid accumulation in microbial cells (Park et al., 2014; Vogt et al., 2014).

**Figure 2.**
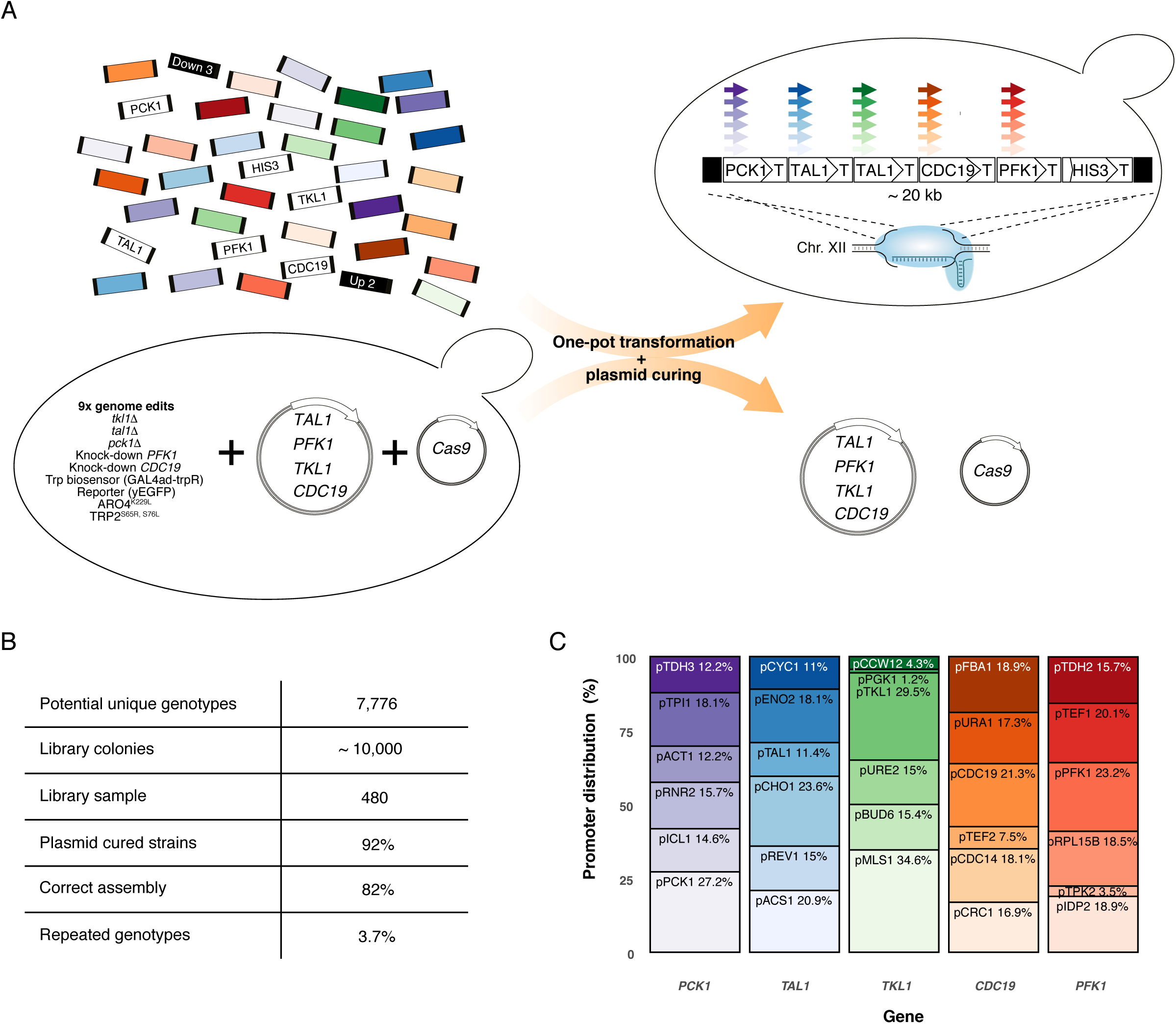
Construction and validation of the 13-parts assembled 20 kb combinatorial promoter:gene library. (A) Strategy for library construction including a 13-part *in vivo* assembly for the reintegration of target genes into a single genomic locus. The platform strain used for one-pot transformation includes a total of 9 genome edits for knowck-out, knock-down and heterologous expression of candidate genes (see METHODS DETAILS). (B) Key descriptive statistics for the library construction and genotyping. (C) Promoter distribution (name, % representation) by gene. Color intensity correlates with promoter strength (see Figure 1D).

### One-pot construction of the combinatorial library

For library construction, we first tested the transformation by constructing five control strains, including a strain with native promoters in front of each of the five selected genes (herein labelled the reference strain; Table S7). Next, we transformed in one-pot the platform strain with equimolar amounts (1 pmol/part) of double-stranded DNA encoding each of the thirty promoters, the five open reading frames encoding the candidate genes with native terminators, a *HIS3* expression cassette for selection, and two 500-bps homology-regions for targeted repair of the genomic integration site. In total, this design combination included 38 different parts for 7,776 unique 20 kb 13-parts assemblies at the targeted genomic locus (Chr. XII, EasyClone site V; Figure 2A). Following transformation, we randomly sampled 480 colonies from the library, together with 27 colonies from the five control strains (507 in total), and successfully cured 423 out of 461 (92%) sufficiently growing strains of the complementation plasmid by means of galactose-induced expression of the dosage-sensitive gene *ACT1* (Figures 2B & S6; Liu et al., 1992; Makanae et al., 2013). Next, genotyping all promoter-gene junctions by sequencing (Figure S2), identified 380 out of 461 (82%) of the sufficiently growing strains to be correctly assembled with only 9 out of 245 (3.7%) of the fully filtered library genotypes observed in duplicates (245 = 250 library and control genotypes - 5 control genotypes)(Figure 2B). Based on a Monte Carlo simulation with 10,000 repeated samplings of 10,000 library colonies, and assuming percent correct assemblies and promoter distribution as determined for the library sample (Figure 2), the expected no. of unique genotypes among all library colonies was calculated to be 3,759. This equals an estimated library coverage of 48% (3,759/7,776). Importantly, all thirty promoters from the one-pot transformation mix were represented in the genotyped designs, with promoters *PGK1* (no. 14) and *MLS1* (no. 15), represented the least (1%) and most (35%), respectively (Figure 2C).

Taken together, these results demonstrate high transformation efficiency of the platform strain, high fidelity of parts assembly, and expected high coverage of the genetically diverse combinatorial library design.

### Engineering a tryptophan biosensor for high-throughput library characterization

In order to support high-throughput analysis of tryptophan accumulation in library strains, we harnessed the power of modular engineering allosterically regulated transcription factors as small-molecule *in vivo* biosensors (Mahr and Frunzke, 2016; Rogers et al., 2016). Here, a yeast tryptophan biosensor was developed based on the *trpR* repressor of the *trp* operon from *E. coli* (Roesser and Yanofsky, 1991; Gunsalus and Yanofsky, 1980). In order to engineer *trpR* as a tryptophan biosensor in yeast, we first tested trpR-mediated transcriptional repression by expressing *trpR* together with a GFP reporter gene under the control of the strong *TEF1* promoter containing a palindromic consensus *trpO* sequence (5’-GTACTAGTT-AACTAGTAC-3’; Yang et al., 1996) downstream of the TATA-like element (TATTTAAG; Figure 3A; Rhee and Pugh, 2012). From this, we observed that *trpR* was able to repress GFP expression by 2.4-fold (Figure S3A). Next, to turn the native *trpR* repressor into an activator with a positively correlated biosensor-tryptophan readout we fused the Gal4 activation domain to the N-terminus of codon-optimized *trpR* (*GAL4_AD_-trpR*) expressed under the control of the weak *REV1* promoter (Figure S3B). For the reporter promoter, we placed *trpO* 97 bp upstream of the TATA-like element of the *TEF1* promoter (Figure S3B), and observed that *trpR* was able to activate GFP expression by a maximum of 1.75-fold upon supplementing tryptophan to the cultivation medium (Figure S3B). To further optimize the dynamic range of the reporter output, the GFP reporter was expressed under a hybrid promoter consisting of tandem repeats of triple *trpO* sequences (i.e., in total 6x *trpO* sequences) located 88 bp upstream of the TATA box in an engineered *GAL1* core promoter without Gal4 binding sites, ultimately enabling *GAL4_AD_-trpR*-mediated biosensing with a dynamic output range of 5-fold, and an operational input range spanning supplemented tryptophan concentrations from ∼2-200 mg/L (Figure 3B).

**Figure 3.**
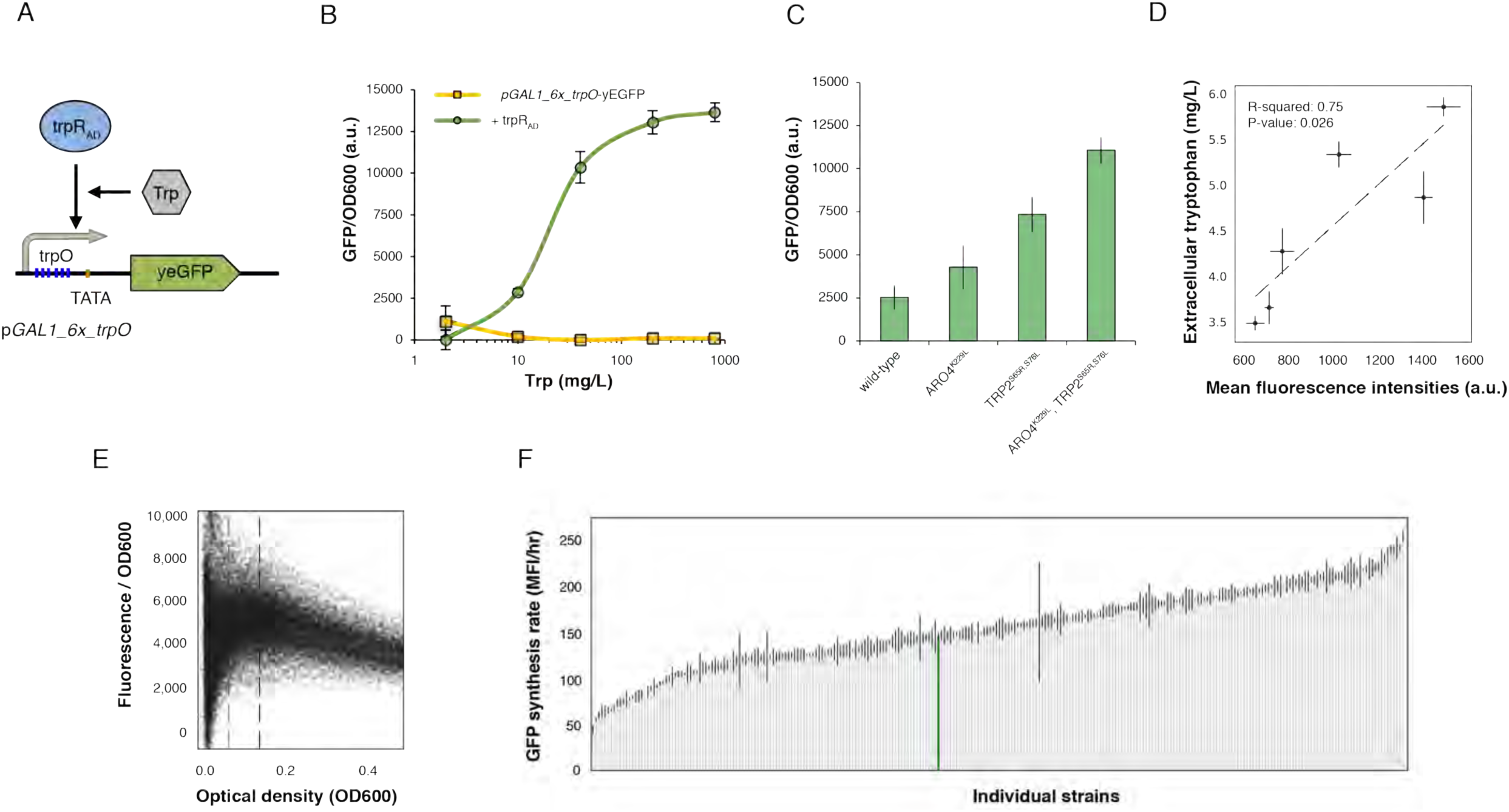
Phenotypic library characterization using an engineered tryptophan biosensor. (A) Schematic illustration of the design of the tryptophan (Trp) biosensor (trpR_AD_) engineered in this study. The trpR_AD_ indicates the engineering tryptophan biosensor comprised of the *E. coli* TrpR fused to the GAL4 activation domain. The biosensor regulates and engineered reporter (yeGFP) *GAL1*-promoter including 6x copies of TrpR binding sites (*trpO*), placed upstream the TATA box of *GAL1* promoter (*pGAL1_6x_trpO*). (B) Fluorescence normalized by optical density (OD600) for two strains related to concentration of tryptophan supplemented media (Mean Fluorescence Intensity/OD, MFI/OD with standard errors, n = 3). Both strains contain the yeGFP reporter under the control of the *pGAL1_6x_trpO* reporter promoter, and only one strain expresses the Gal4 activation domain fused to trpR (in green). (C) Fluorescence normalized by OD600 for a wild-type strain and strains with expression of feedback-resistant versions of ARO4 and TRP2, ARO4^K229L^ and TRP2^S65R,S76L^, respectively (mean fluorescence intensity, MFI/h with standard errors, n = 3). (D) Extracellular tryptophan normalized by OD600 related to fluorescence normalized by OD600 (mean values with standard errors, n = 3). (E) Fluorescence divided by OD600 related to OD600 for library and control strains. Dashed lines are shown at OD600 equals 0.075 and 0.15. (F) Measured mean green fluorescent protein synthesis rate. MFI/h with standard errors, n = 3. The data is ranked according to increasing mean rate. The strain with five native promoters expressing the five candidate genes is highlighted in green. MFI = Mean Fluorescence Intensity. OD600 = Optical density (600 nm). a.u. = arbitrary units.

To further validate the designed biosensor we measured fluorescence output in strains engineered for expression of feedback-resistant versions of ARO4 and TRP2 (ARO4^K229L^ and TRP2^S65R, S76L^; (Hartmann et al., 2003; Graf et al., 1993), and observed high biosensor outputs from these strains in line with previously demonstrated high enzyme activities in strains expressing ARO4^K229L^ and TRP2^S65R, S76L^ (Hartmann et al., 2003; Graf et al., 1993), and thus corroborating the ability of the tryptophan biosensor to monitor changes in endogenously produced tryptophan pools (Figure 3C). Most importantly, we confirmed the biosensor readout as a valid proxy for tryptophan levels, by comparing external tryptophan titers measured by HPLC with a change in GFP intensities for 6 library strains spanning 2.5-fold changes in GFP intensities (*R*^*2*^ = 0.75; Figure 3D).

Having established a biosensor for high-throughput screening of the combinatorial library, we next sought to explore the maximal resolution of the biosensor readout at the single-design level of growing isoclonal strains, with the intention to define optimal data sampling time point. To do so, we measured time-series data of OD and GFP in triplicates for all 507 colonies, covering a total of >144,000 data points (Figure S4). Here, as we observed that the fluorescence per cell generally stabilized at an OD value of 0.075 and started to decrease beyond an OD value of 0.15 (Figure 3E, Figure S4, see METHODS DETAILS), and the between strains variation in fluorescence at the single-cell level was relatively high within this OD-interval, we chose this interval for determining the GFP synthesis rate as a proxy for tryptophan flux. By sampling all variant designs, average GFP synthesis rate was observed to vary between 43.7 and 255.7 MFI/h (approx. 6-fold; Figure 3F), with an average standard error of the mean of 6.6 MFI/h corresponding to an average coefficient of variation for the mean values of 4.3%. By comparison, the GFP synthesis rate of the platform strain, expressing ARO4^K229L^ and TRP2^S65R, S76L^ together with all five candidate genes under native promoters, was 144.8 MFI/h (Figure 3F).

### Using machine learning to predict metabolic pathway designs

Having successfully established a combinatorial genetic library and a large phenotypic data set thereof, we next assessed the potential of using machine learning to predict promoter combinations expected to improve tryptophan productivity. Since there is no algorithm which is optimal for all learning tasks (Wolpert, 1996), we used two different machine learning approaches: the Automated Recommendation Tool (ART) and EVOLVE algorithm (Radivojević et al., 2019; TeselaGen, 2019). The input for both algorithms was the promoter combination and tryptophan productivity (measured through the GFP proxy, Figure S4). Briefly, ART uses a Bayesian ensemble approach where eight regressors from the scikit-learn library (Pedregosa et al., 2011) are allowed to “vote” on a prediction with a weight proportional to their accuracy; the EVOLVE algorithm is inspired by Bayesian Optimization and uses an ensemble of estimators as a surrogate model that predicts the outcome of the process to be optimized (see METHODS DETAILS). As the quality of the data is of paramount importance for machine learning predictions, we initially filtered our data to avoid genotypes with insufficient growth, no sequencing data, incorrect assembly, no plasmid curation, or which exhibited more than one genotype (see METHOD DETAILS; Figure S5). Following this, approximately 58% (266/461) of the growing strains remained after filtering, while another 3% of the remaining data was removed because of lack of reproducibility (high error in triplicate measurements)(Figure S5).

Both modeling approaches, ART and EVOLVE, were able to recapitulate the data they were trained on. The average (obtained from 10 independent runs) training mean absolute error (MAE) of the predicted tryptophan production compared to the measured values was 13.8 and 11.9 MFI/h for the ART and EVOLVE model approaches, respectively, when calculated for the whole data set (Figure 4A-B). These MAEs represent ∼7% and 6% of the full range of measurements (50 to 200 MFl/h). The train MAE uncertainty (represented by the shaded area in Figure 4A-B and quantified as the 95% confidence interval from 10 runs) decreased slightly with increasing size of the training data set for ART, whereas the overall uncertainty was smaller for the EVOLVE model approach (Figure 4A-B). The ability to predict the production for new promoter combinations the algorithms had not been trained on was tested by cross-validation, i.e. by training the model on 90% of the data, and then testing the predictions of this model against measurements for the remaining 10% (10-fold cross-validation). Here, the average cross-validated MAE (test MAE) was 21.4 and 22.4 MFI/h for ART and EVOLVE model approaches, respectively (Figure 4A-B), which represent ∼11% of the full range of measurements. The test MAE decreased systematically with the size of the data set, yet the decrease rate declined markedly as more data was added. However, while the two approaches had similar average cross-validated MAEs, the uncertainty of the MAEs was slightly smaller for ART than for EVOLVE algorithm (Figure 4A-B).

**Figure 4.**
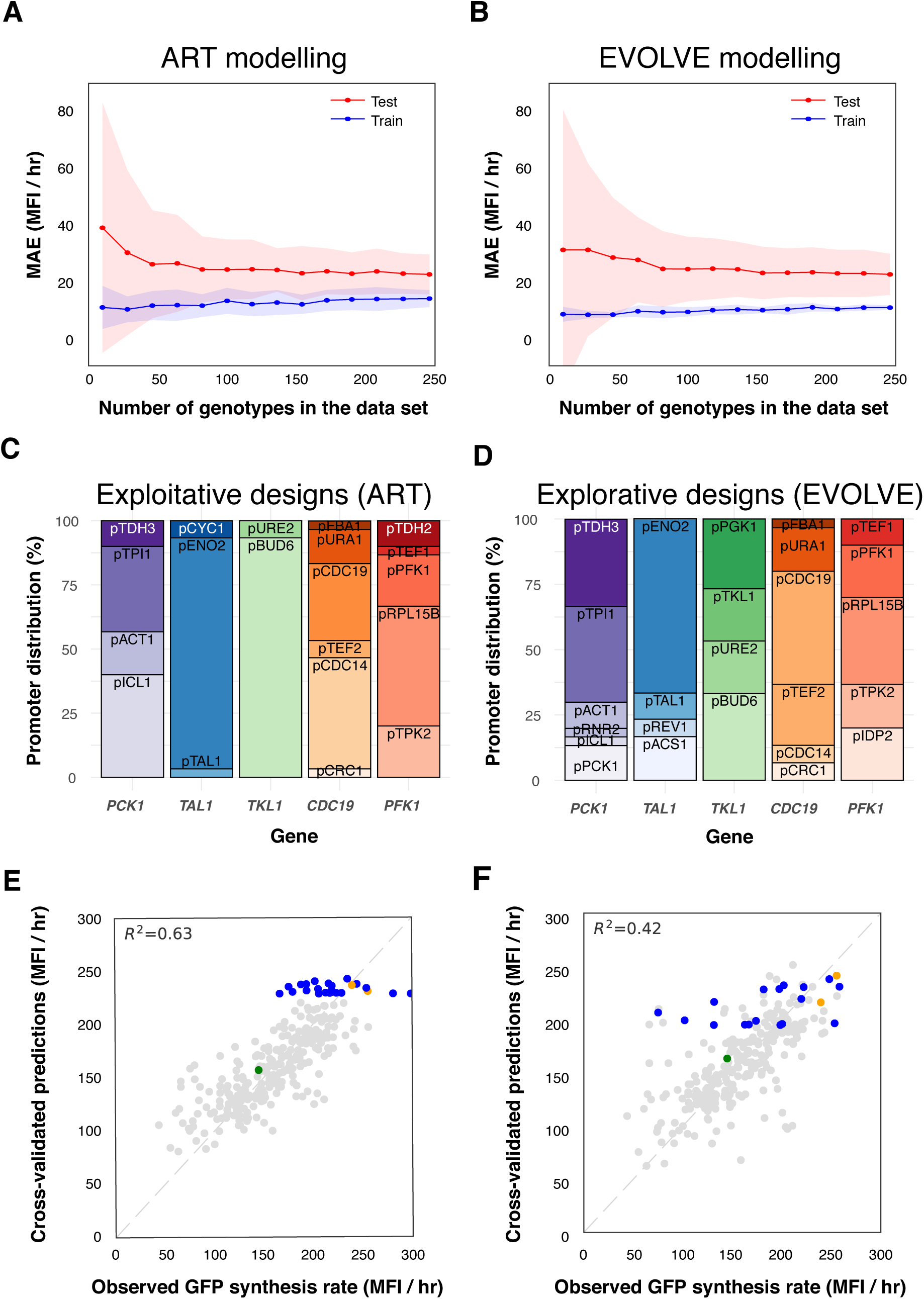
Machine learning-guided predictive engineering of tryptophan metabolism. (A-B) Learning curves for ART and EVOLVE algorithms, respectively. Mean absolute error (MAE) from model training and testing as a function of the number of genotypes in the dataset. Shaded areas represent 95% confidence intervals. Blue curves indicate MAE when calculated for the whole data set (Train), while red curves indicate the cross-validation, i.e. by training the models on 80% of the data and then testing the predictions of this model against measurements for the remaining 20% (Test). (C-D) Promoter distributions for the 30 recommendations of the exploitative (ART) and explorative (EVOLVE) approach, respectively. The orders and colors of promoters correspond to those in Figure 1C. (E-F) Cross-validated predictions vs average of measured GFP synthesis rate for the exploitative (ART) and explorative (EVOLVE) approach, respectively. Data is shown for library and controls strains (grey markers; green markers show the platform strain expressing ARO4^K229L^ and TRP2^S65R,S76L^), as well as for recommended strains (blue markers; orange markers show recommendations that overlap between the two approaches).

### Machine learning-guided engineering of designs with high tryptophan productivity

Next, beyond enabling prediction of tryptophan production, we used an exploitative approach implemented in the ART model and an explorative one adopting the EVOLVE algorithm to recommend two sets of 30 prioritized designs aiming for high tryptophan production (<u>Tables S8 and S</u>9). The exploitative model focuses on exploiting the predictive power to recommend promoter combinations that improve production, whereas the exploratory model combines predictive power with the estimated uncertainty of each prediction, to recommend promoter combinations (Radivojević et al., 2019; TeselaGen, 2019).

Among the recommendations from each of the two machine learning approaches, two overlapped (SP588 and SP627, Table S8-S9). Interestingly, while use of *PGK1* promoter to control *TKL1* expression was underrepresented in the original library sample (Figure 2C), the explorative set of recommendations included eight (even top-three) designs with *PGK1* promoter for expression control of *TKL1*, and the exploitative approach included none (Table S5; Figure 4C-D). From construction of these recommendations, we used the same genome engineering approach as for library construction (Figure 2A) to successfully construct 19 individual assemblies of the explorative recommendations and 24 individual assemblies of the exploitative recommendations. Interestingly, we were not able to construct any of the eight designs with *PGK1* promoter, partially explaining the lower number of viable strains found with the explorative approach.

Of the 41 recommendations constructed, the predictions from both sets generally fitted well with the measurements, and both approaches successfully enabled predictive strain engineering for high-performing GFP synthesis rates, with the best recommendation having a measured GFP synthesis rate 106% higher than the already improved platform design, and 17% higher than the best one in the library sample (Figure 4E-F). Moreover, eight recommendations were found in the top-ten of productivity, of which four were from the exploitative set, three were from the explorative set, and one overlapping between the two sets. Comparing the output of the ART and EVOLVE approaches, the variation in measurements was higher for strains recommended with the explorative EVOLVE approach than for strains recommended with the exploitative ART approach (Figure 4E-F), and the explorative approach included recommendations based on a more diverse set of promoters than the exploitative approach (Figure 4C-D). Still, taken together, both approaches successfully enabled predictive engineering of a strain with tryptophan productivity beyond those previously observed (Figure 4E-F).

## DISCUSSION

We have demonstrated that mechanistic and machine learning approaches can complement and enhance each other, enabling a more effective predictive engineering of living systems. Using a single design-build-test-learn cycle, this study i) leveraged mechanistic genome-scale models to select and rank reactions/genes most likely to affect production, ii) included the efficient one-pot construction of a library with different promoter combinations for these reactions, and iii) used machine learning algorithms trained on the ensuing phenotyping data to choose novel promoter combinations that further enhance tryptophan productivity. In total, we managed to increase the tryptophan synthesis rate by 106% compared to an already improved reference strain (ARO4^K229L^ and TRP2 ^S65R, S76L^).

To gather the large data sets required to enable machine learning approaches, we developed a biosensor which enabled the sampling of >144,000 GFP intensity measurements as a proxy for tryptophan flux for 1,728 isoclonal designs in a high-throughput fashion (Figures 3E, S5A). Indeed, while requiring a few design iterations (Figures 3A, S3), the tryptophan biosensor ultimately allowed us to i) phenotypically characterize an order of magnitude higher number of strains than in previous machine learning-guided metabolic engineering studies (Alonso-Gutierrez et al., 2015; Lee et al., 2013a; Redding-Johanson et al., 2011; Zhou et al., 2018a), and ii) identify optimal sampling points that displayed the largest differences between genotypes (Figures 3C, S4). Likewise, one-pot CRISPR/Cas9-mediated genome editing was a vital enabling technology for this project, since it allowed us to efficiently create a diverse 20-kb clustered combinatorial library with representation of all 30 specified sequence- and expression-diverse promoters to control five expression units, including very few duplicate designs (Figure 2B-C).

Enabled by this high-quality data set, we used two different machine learning models for predicting productivity (ART and EVOLVE algorithm), and two different approaches to recommend new strains (exploitative and explorative). Cross-validation showed that both models could be trained to show good correlations (MAE approximately 11% of the measurement range) between predictions and measurements for data they had not seen previously (test data). The test MAE was basically the same for the two models, and plateaued quickly as a function of the number of genotypes in the training data set (Figure 4A-B). Whereas the uncertainty in predictive accuracy decreased considerably with the number of genotypes in the data set, this decrease was similar for both models. With this in mind, a relevant guideline for choosing a recommendation approach should focus on the desired outcome: the explorative approach providing a more diverse set of recommendations (Figure 4C-D), whereas the exploitative approach provides less varied recommendations. We observed the largest improvement in productivity when using the exploitative approach (Figure 4E-F). However, if subsequent design-build-test-learn cycles are performed, the diversity of recommendations of the explorative approach could help avoid local optima of tryptophan production(Figure 4E-F).

Notably, while the recommendations were able to improve production, the predictions from both machine learning models were noticeably worse than for the library, reflecting the general challenge of extrapolating outside of the previous range of measurements. As such, we envision that future machine learning approaches will need to focus on models able to extrapolate more efficiently.

With respect to advancing biological understanding of tryptophan metabolism, the results provided examples of anticipated results as well as non-intuitive predictions. The best performing strain (SP606, Table S8) predicted by machine-learning, displayed knock-downs of both *CDC19* and *PFK1*, corroborating our intuitive strategies for increasing precursor availability: i.e. lower pyruvate kinase activity would lead to higher PEP pools, while limiting glycolysis redirects carbon flux into PPP and subsequently increases E4P. However, this strain also had low expression of *TKL1* and high expression of *TAL1*, despite the report that overexpression of *TKL1*, rather than *TAL1*, leads to higher aromatic amino acid production in both *E. coli* and yeast (Curran et al., 2013). This finding remarks the importance of carefully considering the systems-level context of these “metabolic rules of thumb” (e.g. overexpress TKL1 instead of TAL1 for higher amino acid production) to ensure their validity. Consistently, both the second (SP616) and third (SP624) best performing strains, also predicted by machine learning, had low expression of *TKL1* and high expression of *TAL1*, together with very low expression (*TPK2* promoter) for *PFK1* and high expression of *CDC19*. One possible explanation is that, although normally expressed, the pyruvate kinase activity could be limited by low level of its allosteric activator FBP due to limited PFK expression. Another plausible explanation is that medium-high expression of *PCK1* (conversion of oxaloacetate to PEP) by *ACT1* or *TDH3* promoters in these two strains can replenish PEP pools consumed by pyruvate kinase. The fact that 8 out of 10 top-performing strains had high expression of *PCK1*, which was not predicted to be impactful on glucose by the GSM approach, indicates that this indeed has a positive effect on tryptophan biosynthesis rate, and stresses the importance of combining mechanistic and machine learning approaches.

Ultimately, in our case study, machine learning models have demonstrated significant predictive power. However, this predictive power is heavily dependent on the availability of high quality experimental data, which is not a prerequisite for mechanistic GSMs. Without any experimental input, GSMs are able to guide metabolic engineering using various constraint-based algorithms, which, however, predict a large number of potential targets and may also miss some effective ones, e.g. *PFK1* in our study. This could be due to the lack of other information beyond metabolism e.g. regulation in GSMs. To address this problem, manual efforts are currently needed to filter out less relevant targets, and add intuitively promising ones based on existing knowledge and literature mining. Additionally, future GSMs that include more biological aspects and suitable predicting algorithms are envisioned to further improve gene target selection. Irrespective of the ongoing efforts for model-guided engineering of living cells, this study highlights the enhanced predictive power obtained by combining GSMs for selecting genetic targets with machine learning algorithms for leveraging experimental data. Finally, as even more efficient methods for combining data-driven machine learning algorithms and GSMs are developed, we envision dramatic improvements in our ability to engineer virtually any cell system effectively.

## ACKNOWLEDGMENTS

This work was supported by the Novo Nordisk Foundation and the European Commission Horizon 2020 programme (grant agreement No. 722287 and No. 686070). This work was also part of the DOE Agile BioFoundry (http://agilebiofoundry.org), supported by the U.S. Department of Energy, Energy Efficiency and Renewable Energy, Bioenergy Technologies Office, and the DOE Joint BioEnergy Institute (http://www.jbei.org), supported by the Office of Science, Office of Biological and Environmental Research, through contract DE-AC02-05CH11231 between Lawrence Berkeley National Laboratory and the U.S. Department of Energy. The Department of Energy will provide public access to these results of federally sponsored research in accordance with the DOE Public Access Plan (http://energy.gov/downloads/doe-public-access-plan). H.G.M. was also supported by the Basque Government through the BERC 2014-2017 program and by Spanish Ministry of Economy and Competitiveness MINECO: BCAM Severo Ochoa excellence accreditation SEV-2013-0323. This work was also supported by the Chilean economic development agency, Corfo, through grant 17IEAT-73382.

## AUTHOR CONTRIBUTIONS

JZ, SDP, JDK, JN and MKJ conceived the study. JZ and SDP conducted all experimental work, YC and BJS all mechanistic modelling, and TR, ZC, and HGM developed and applied statistical modelling and recommendations based on ART, while EA, AR, and MF developed and applied statistical modelling and recommendations based on TeselaGen EVOLVE model. SDP, JZ, and MKJ wrote the manuscript.

## TABLES

### STAR*METHODS

Detailed methods are provided in the online version of this paper and include the following:

- KEY RESOURCES TABLE
- CONTACT FOR REAGENT AND RESOURCE SHARING
- EXPERIMENTAL MODEL AND SUBJECT DETAILS
- METHOD DETAILS
  - Mechanistic modeling of high tryptophan flux
  - Promoter selection
  - General strain construction
  - Platform strain construction
  - Construction of combinatorial library
  - Development of tryptophan biosensor
  - Validation of biosensor by HPLC
  - Genomic DNA sequencing
  - Measuring fluorescence and growth
- QUANTIFICATION AND STATISTICAL ANALYSIS
  - Modelling
- DATA AND SOFTWARE AVAILABILITY

## STAR*METHODS

Detailed methods are provided in the online version of this paper and include the following:

### KEY RESOURCES TABLE

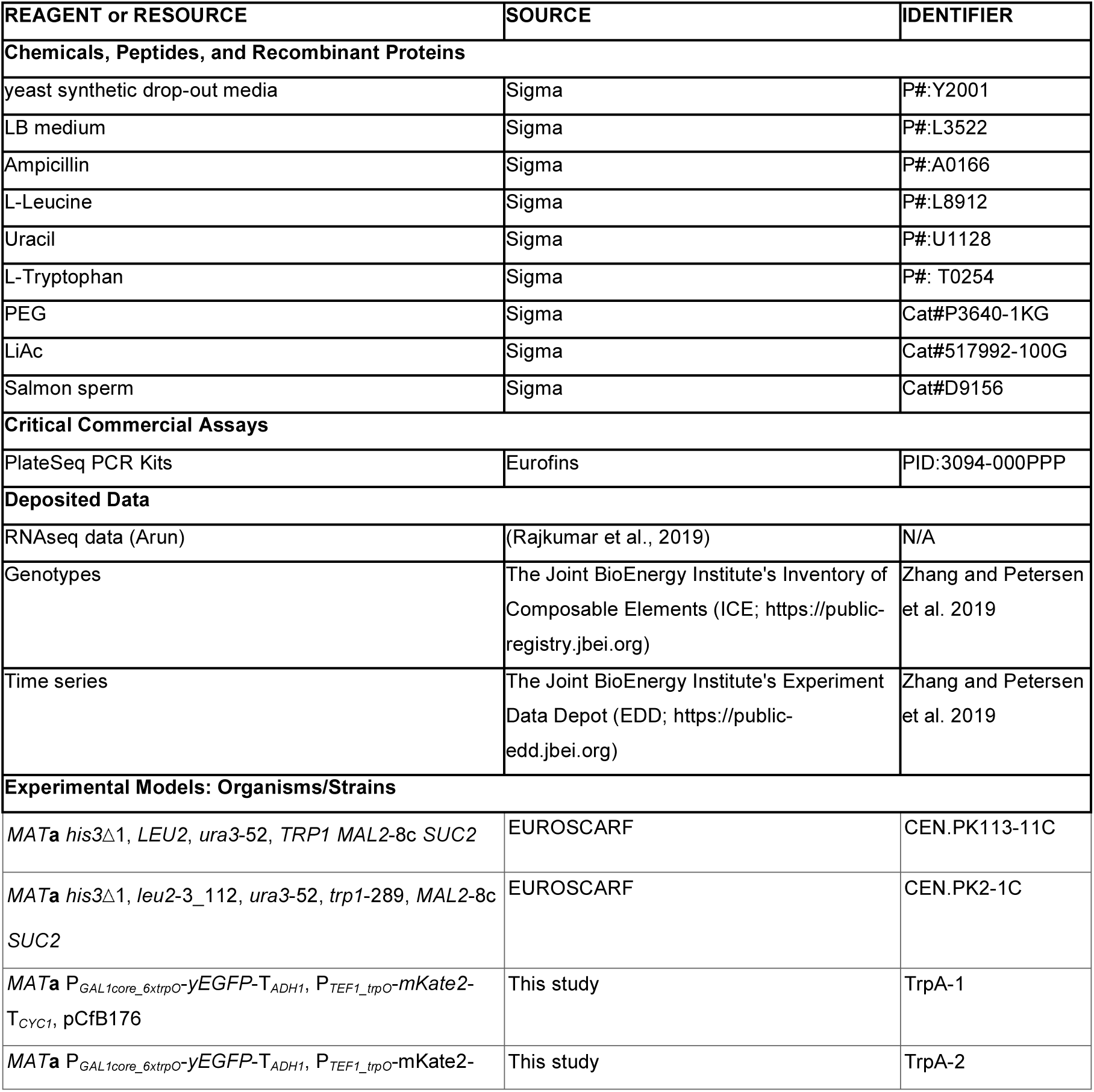
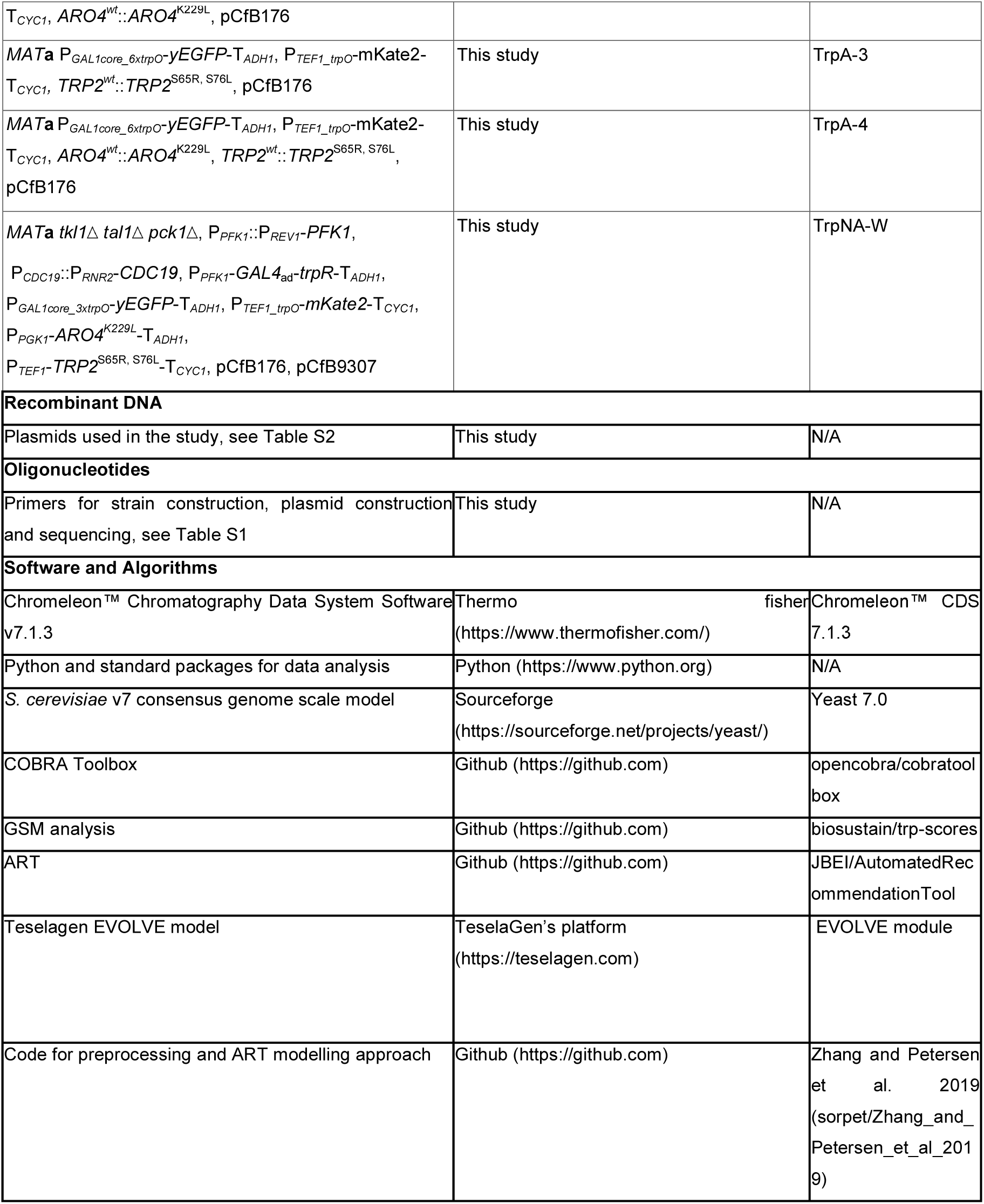

## CONTACT FOR REAGENT AND RESOURCE SHARING

Further information and requests for resources and reagents should be directed to and will be fulfilled by the Lead Contact, Michael Krogh Jensen.

## EXPERIMENTAL MODEL AND SUBJECT DETAILS

*Saccharomyces cerevisiae* strains were derived from CEN.PK2-1C (EUROSCARF, Germany). These were cultivated in yeast synthetic drop-out media (Sigma-Aldrich) at 30 °C. *Escherichia coli* DH5α were cultivated in LB medium containing 100 mg/l ampicillin (Sigma-Aldrich) at 37 °C.

## METHOD DETAILS

### Mechanistic modeling of high tryptophan flux

In order to select targets for increased tryptophan accumulation, we followed a constraint-based strategy implemented in a recent study (Ferreira et al., 2019), similar to the FSEOF approach (Choi et al., 2010). Briefly, flux balance analysis (FBA; Orth et al., 2010) was used to simulate growth of *S. cerevisiae* at 11 different sub-optimal growth conditions ranging from 30% to 80% of the maximum specific growth rate, with all remaining flux oriented towards tryptophan accumulation. Based on these simulations, a score was calculated for each reaction in metabolism as the average simulated flux fold-change compared to maximum growth rate conditions. These reaction scores were in turn used to compute gene scores, by averaging the associated reaction scores. A gene score higher than one means that the gene is associated to reactions that increase in flux as tryptophan production increases, and could point to a target for overexpression. On the other hand, a gene score lower than one signifies that the gene is connected to reactions that decrease their flux as tryptophan production increases, and therefore could be a target for downregulation. The analysis was performed with either glucose or ethanol as carbon sources, so to find candidates under a mixed-fermentation regime, a purely respiratory regime and the overlap between both regimes. The 7th version of the consensus genome-scale model of *S. cerevisiae* (Aung et al., 2013), a parsimonious FBA (pFBA) approach (Lewis et al., 2010), and the COBRA toolbox (Heirendt et al., 2019) were used for all simulations.

### Promoter selection

Each of the five gene targets was expressed under six unique promoters. The six promoters included the promoter native to the gene as well as 5 promoters chosen to span a wide expression range All promoters were chosen based on absolute mRNA abundances measured for S. cerevisiae CEN.PK 113-7D in the mid-log phase (Rajkumar et al., 2019), and unless otherwise stated were 1 kb in length by default. To minimize homologous recombination during one-pot transformation for library construction and potential loop-out of promoters and genes following genomic integration, all scanned promoter sequences were aligned to ensure there were no extensive homologous sequence stretches.

### General strain construction

Strains were edited using the CasEMBLR method (Jakočiu□nas et al., 2015). All integration were directed towards EasyClone sites (Jensen et al., 2014). Homology regions between DNA parts were by default 30 bp, and homology regions, framing the repair assembly, were about 0.5 kb. Yeast transformations were performed by LiAc/SS carrier DNA/PEG method (Gietz and Schiestl, 2007). DNA parts and plasmids were purified using kits from Macherey-Nagel. PCR products for USER assembly were amplified using Phusion U Hot Start PCR Master Mix (ThermoFisher), bricks for transformation by Phusion High-Fidelity PCR Master Mix with HF Buffer (ThermoFisher), whereas colony PCRs were performed using 2xOneTaq Quick-Load Master Mix with Standard Buffer (New England Biolabs). Genomic DNA was extracted from overnight cultures using Yeast DNA Extraction Kit (Thermo Scientific). Oligos were purchased from IDT. Sequencing was performed by Eurofins. All primers, plasmids, and yeast strains, are listed in <u>Tables S1, S2, and S3</u>, respectively.

### Platform strain construction

Several enzymes within the aromatic amino acid (AAA) biosynthesis are subject to allosteric regulations. Specifically, 3-deoxy-D-arabino-heptulosonate-7-phosphate (DAHP) synthase (encoded by *ARO4*), which controls the entry of the shikimate pathway, is feedback inhibited by all three aromatic amino acids, although to different extents. Anthranilate synthase (encoded by *TRP2*), which catalyzes the first committed step towards the tryptophan branch, is also inhibited by its end product tryptophan (Braus, 1991). To maximise the transcriptional regulatory effect on the tryptophan flux, and benchmark with current state-of-the-art in shikimate pathway optimization, feedback resistant variants of these two enzymes, ARO4^K229L^ (Hartmann et al., 2003) and TRP2^S65R, S76L^ (Graf et al., 1993), were overexpressed under the *TEF1* and *TDH3* promoters, respectively at EasyClone site XI-3 (Jessop□Fabre et al., 2016; Table S2). Secondly, a tryptophan biosensor system (see Library phenotypic characterization) was introduced by integrating corresponding sensor and reporter sequences into EasyClone sites at Chr. XI-2 and XI-5, respectively (Jensen et al., 2014).

### Construction of combinatorial library

Due to the dramatic decrease in transformation efficiency targeting multiple loci in the genome (Jakočiūnas et al., 2015), we opted for removing all five target genes from their original loci and assemble the five expression units into a single cluster for targeted integration into EasyClone site XII-5 (Jensen et al., 2014), and thereby ensuring comparable genomic accessibility of all genes. While *PCK1, TKL1* and *TAL1* were successfully knocked out; deleting *PFK1* and/or *CDC19* was unsuccessful. Alternatively, we replaced *PFK1* and *CDC19* promoters with weak *REV1* and *RNR2* promoters, respectively. Due to an expected loss of activity in phosphofructokinase (PFK1) and pyruvate kinase (CDC19), and consequently slow ATP generation, the resulting strain (TrpNA-W) grew extremely poorly and was barely transformable using linear DNA fragments for assembly. To overcome this limitation, the TrpNA-W strain was complemented with plasmid pCfB9307 (Table S2) harboring *PFK1, CDC19, TKL1* and *TAL1* genes, which restored the growth to the wild type level. The plasmid backbone carries yeast *ACT1* gene under the control of *GAL1* promoter, which can be used as counter-selection of the plasmid due to the growth arrest caused by *ACT1* overexpression on galactose as the sole carbon source (Makanae et al., 2013, Figure S6).

For combinatorial library construction we adopted CasEMBLR (Jakočiu□nas et al., 2015). Briefly, five target genes together with a *HIS3* expression cassette (in the order of *PCK1-TAL1-TKL1-CDC19-PFK1-HIS3*) were assembled in the same orientation and integrated at EasyClone site XII-5 (Jensen et al., 2014). All five target genes (the complete ORFs) together with their terminators (500 bp downstream of the stop codon) were amplified from the genomic DNA of yeast strain CEN.PK113-7D using primers listed in Table S1. All 30 promoters (defined as the 1000 bp upstream the ORF) were amplified using primers with a 30 bp overlap to adjacent DNA parts (i.e. the terminator upstream and the target gene). All promoters can be found in <u>Tables S4</u>. The *HIS3* cassette was amplified from plasmid pRS413-HIS3 (Sikorski and Hieter, 1989) with primers 30 bp overlapping with the *PFK1* terminator and fragment homologous to the downstream of XII-5. The *HIS3* cassette was included as one part of the assembly. The one-pot transformation of all 38 parts (30 promoters, 5 candidate genes, *HIS3* cassette, and up- and down-homology regions for EasyClone site XII-5) was performed with 50 mL the base strain grown to an optical density of 1.0 (equivalent to 6.5 mg of cell dry weight), 5.0 ug of plasmid expressing the guide RNA targeting XII-5, and 1.0 picomole of each of 13 DNA fragments. A total of 480 colonies were picked from 10 transformation plates by dividing the area of each individual plate into 4 subareas of equal size and picking 12 colonies of varying size from each subarea.

Finally, the complementation plasmid introduced was cured by culturing strains to stationary phase twice in media with galactose instead of glucose as carbon source (Figure S6). The success of curing were then gauged by a growth assay where LEU auxotrophs were considered as cured and prototrophs as not cured. Control strains and recommended strains were constructed similarly to the library strains except that instead of transforming pools of promoter parts for each gene only specific promoters were transformed per gene.

### Development of tryptophan biosensor

The yeast tryptophan biosensor was developed based on the *trpR* repressor of the *trp* operon from *E. coli* (Gunsalus and Yanofsky, 1980). The *trpR* gene was amplified from E. coli M1665 genome. All yeast promoters as well as the activator domain of *GAL4* were amplified from *S. cerevisiae* strain CEN.PK113-7D genome. All designs of trpR biosensor and GFP reporter were first cloned into the pRS416 (*URA3*) and pRS413 (*HIS3*) vectors, respectively, by USER cloning (Bitinaite et al., 2007). The activator domain of *GAL4* (*GAL4*_AD_) was fused to trpR with a GSGSGS linker by USER cloning. The *trpO* sequence was inserted into the *TEF1* promoter 8 bp downstream of the TATA-like element (TATTTAAG) by inverse PCR from a plasmid containing the P_*TEF1*_-*yEGFP-T*_*ADH1*_ cassette, with both primers containing the overhang AACTAGTAC (ie., half of the *trpO* sequence). The linear PCR product was treated with DpnI enzyme to fragmente the template plasmid and self-ligated to generate circular plasmid (Quick Ligation™ Kit, NEB). Promoters containing multiple *trpO* sequences were constructed by USER cloning from a synthetic DNA fragment (Integrated DNA Technologies) of a minimal *GAL1* promoter (−329 to −5 relative to the *GAL1* open reading frame, thus without the *GAL4* binding sequence which is located at −435 to −418) with 3x tandem repeats of *trpO* (separated by 2 nucleotides) inserted at 88 bp upstream of the TATA box (TATATAAA). Plasmids containing the sensor and reporter cassettes were transformed into yeast strain CEN.PK113-11C. To test the biosensor performance, yeast transformants were grown in selection media overnight and regrown in Delft medium supplemented with various tryptophan concentrations (2-1000 mg/L) for 6 hrs (typically reaching early exponential phase). GFP and mKate2 outputs were measured on SynergyMX microtiter plate reader (BioTek) with excitation/emission at 485/515 nm and 588/633 nm, respectively, and always normalized by absorbance at 600 nm (OD600nm). To construct the base strain for library assembly, the tryptophan sensor (P_*REV1*_-*GAL4*_*AD*_-*trpR-T*_*ADH1*_) and the reporter cassette (P_*GAL1core_3xtrpO*_-*yEGFP*-T_*ADH1*_xs, P_*TEF1_trpO*_-*mKate2*-T_*CYC1*_) were integrated into strain TC-3 (Jakočiūnas et al., 2015) at the EasyClone sites XI-2 and XI-5 (Jessop□Fabre et al., 2016), respectively.

### Validation of biosensor by HPLC

To validate the correlation between biosensor reporter gene output and tryptophan production, we quantified extracellular tryptophan levels by HPLC using a method described by Luo et al. (2019). Supernatants of cultivated strains were separated from the culture broth following 24 hrs of cultivation in synthetic dropout medium without tryptophan and histidine. From this 200 µl was used for HPLC and the data were processed using Chromeleon™ Chromatography Data System Software v7.1.3.

### Genomic DNA sequencing

Genomic DNA was extracted from overnight cultures using method described by Lõoke et al. (2011). Each extract was used as template in 5 PCR reactions spanning the 5 integrated promoters and amplifying from 1,200 - 1,700 bp. The PCR products were validated using a LabChip GX II (Perkin Elmer) and sequenced using PlateSeq PCR Kits (Eurofins) according to the manufacturer’s instructions. From the LabChip results, a PCR reaction was considered as trusted if it showed a strong band of the correct size, not trusted if it showed a strong band of the wrong size, and as no information gained if it showed a weak or no band. From the sequencing results, a sequencing reaction was considered as trusted if it showed an unambiguous sequence of the expected length (i.e. only limited by length of PCR fragment, stretches of the same nucleotide in the promoter or of about 1,000 bp limit of sanger sequencing reactions), not trusted if it showed an unambiguous sequence of the expected length with an assembly error, and no information gained if there were no or bad sequence results. If one or more sequencing results from the same strain showed double peaks in the promoter region the strain was considered as a double population. Finally, the promoter was noted as failed assembly (FA) if either LabChip and or sequencing results were considered not trusted, as no information (NI) if the sequencing result was no information and else as the promoter predicted by pairwise alignment between sequencing results and promoter sequence.

### Measuring fluorescence and growth

Yeast cells were cultured ON to saturation, diluted to OD_600_ 0.025 (measured by reading the absorbance at 600 nm on Synergy Mx Microplate Reader, BioTek) and then cultured again in a Synergy Mx Microplate Reader. While culturing, the reader measured OD_600_ and fluorescence with excitation and emission wavelengths of 485 and 515 nm, respectively every 15 min for 20 hrs. All wells were sealed with VIEWseal membrane (Greiner Bio-One).

## QUANTIFICATION AND STATISTICAL ANALYSIS

### Modelling

All genotype and time series data as well as scripts for preprocessing are publicly available (see section DATA AND SOFTWARE AVAILABILITY). Briefly, all OD and GFP measurements were subtracted background signal (i.e. mean value of OD and GFP measurements in wells containing pure media). Background signals were calculated for each 96-well plate. Strains were quality-controlled based on 5 criteria. The criteria were: 1. Optical densities must cover the whole range up to 0.15 OD units to exclude uninoculated wells and wells with insufficient growth, 2. Sequencing results must exist for all five promoter gene junctions, 3. The integrated sequence must be exactly as designed, 4. The complementation plasmid must be cured, and 5. The sequencing results must not indicate the presence of multiple genotypes (Figure S5A). GFP synthesis rates were calculated in the OD_600_ interval from 0.075 to 0.150, as measured by a Synergy Mx Microplate Reader from BioTek.

In the ART approach, outliers were identified and removed based on replicate differences in GFP synthesis rate relative to the mean value for the strain. Replicates with the one percent most extreme differences were identified and the corresponding strains were removed. GFP synthesis rate was modelled as a function of promoter combination, represented through one-hot encoding, using the Automated Recommendation Tool (ART; Radivojević et al., 2019). Briefly, ART uses a probabilistic ensemble model consisting of eight individual models. The weight of each ensemble model is considered a random variable with a probability distribution characterized by the available training data, and determined through Bayesian inference and Markov Chain Monte Carlo (Brooks et al., 2011). ART uses the trained ensemble model in combination with a Parallel Tempering approach (Earl and Deem, 2005) to recommend 30 new promoter combinations (unseen designs), which are predicted to improve production. The recommended designs were chosen as the 30 strains with the highest expected GFP synthesis rate predicted by the model. This recommendation approach was labelled exploitative since predictions with high uncertainty were not prioritized, although ART can provide both exploitative and explorative recommendations

For the TeselaGen EVOLVE algorithm used in this study, outliers were identified and removed based on a method described by Rousseeuw and Hubert (2011). The decision was made on a per strain basis taking into account replicate to mean value differences. In cases where just a single replicate was left after filtering, this replicate were excluded as well. Of the remaining strains, GFP synthesis rate were modelled as a function of promoter combination coded as categorical variables using a TeselaGen-developed machine learning algorithm based on Bayesian Optimization (Mockus, 1994). The algorithm was set-up to recommend 30 new promoter combinations (unseen designs), and designs were chosen by highest selection score. The selection score was the expected improvement (Bergstra et al., 2011), calculated based on predicted high GFP synthesis rate and the uncertainty of prediction. The approach was labelled explorative since high uncertainty weighed positively in the selection score calculation. While using EVOLVE for explorative recommendations, thereby complementing the ART approach, it should be mentioned that EVOLVE can be set up to provide both explorative and exploitative recommendations.

## DATA AND SOFTWARE AVAILABILITY

The complete flux balance analysis, with additional simulation details and filtering criteria, is publicly available at https://github.com/biosustain/trp-scores. The genotype and time series datasets generated during this study are available at The Joint BioEnergy Institute’s Inventory of Composable Elements (ICE; https://public-registry.jbei.org) and Experiment Data Depot (EDD; https://public-edd.jbei.org), respectively under the study ‘Zhang and Petersen, et al 2019’ (Ham et al., 2012; Morrell et al., 2017). The complete preprocessing and all statistical calculations are documented in a jupyter notebook, available at https://github.com/sorpet/Zhang_and_Petersen_et_al_2019. The notebook also contains the ART approach for modeling and strain recommendations. The Teselagen software is available through commercial and non-commercial licenses (https://teselagen.com).

## SUPPLEMENTAL ITEM TITLES

**Figure S1. Related to Figure 1. Dendrogram of the sequence diversity of 30 selected native yeast promoters**. Sequence pTEF1c1a with a single nucleotide change from pTEF1 has been added as a reference. The dendrogram was constructed using the neighbor-joining method (Saitou and Nei, 1987; Studier and Keppler, 1988).

**Figure S2. Related to Figure 1. Genotyping strategy**. Schematic outline of the genotyping strategy to assess correct *in vivo* junction-junction assemblies of 11 parts, and the integration at EasyClone site XII-5 (Jensen et al., 2014). Marked in red are chromosomal regions of EasyClone site XII-5, whereas green marks the promoters, and yellow the coding sequences and terminators. Marked in blue is the selectable *HIS3* expression cassette, while genotyping PCRs are marked in light red. Primers used for sequencing of the 5 PCR reactions are marked seq1-seq5.

**Figure S3. Related to Figure 3. Biosensor development and characterization**. Overnight cultures of the strain containing sensor and reporter was used to inoculate fresh media supplemented with various concentrations of tryptophan and grown for 6 hours (early-mid exponential phase). Optical density (measured as absorbance at 600 nm) was used to normalize the green fluorescence (excitation/emission at 485/515 nm). (A) *E. coli trpR* was directly expressed in a yeast strain harboring the yEGFP reporter under the control of *TEF1* promoter containing *trpO* sequence inserted downstream of the TATA-like element. (B) The *trpR* gene was fused to the C-terminus of the activator domain of GAL4 (GAL4_ad_) with a GSGSGS linker, turning this transcriptional repressor into an activator (trpAD). Accordingly, the *trpO* sequence was placed upstream of a truncated *TEF1* promoter (lacking region with multiple Rap1-binding sites).

**Figure S4. Related to Figure 3E-F. Parameter estimation from time series data**. (A) Representative growth curve of *S. cerevisiae* in microtiter plates. *S. cerevisiae* was grown in yeast synthetic drop-out media in 96-well microtiter plates, and cell density measured at 600 nm (OD_600_) over 24 hrs. (B) Representative tryptophan biosensor output measured as fluorescence (GFP) in *S. cerevisiae* cells (n = 1). *S. cerevisiae* was grown in yeast synthetic drop-out media in 96-well microtiter plates, and GFP measured at 485 nm (OD_485_) over 24 hrs. (C) Tryptophan biosensor output normalized by absorbance at 600 nm (OD_600_) over 24 hrs. For (A-C) the red line shows model fitting using a univariate spline. All plots represent a single replicate measurement (n = 1). The green, yellow and blue markers indicate OD_600_ = 0.075, OD_600_ = 0.15, and maximum rate of OD_600_ increase, respectively.

**Figure S5. Related to Figures 3-4. Data filtering and outlier removal**. (A) Schematic illustration of the various filtering steps applied for data quality control. The six steps used for filtering are indicated by number to the left, and listed to the right are the numbers of unique genotypes as inferred from sequencing, the number of strains, and the number of experimental units (Exp. units, n = 3). (B) The distribution of absolute differences between replicate measurements (n = 3) of strain GFP synthesis rate. (C) Same as in (B), but with y-axis expanded by a factor 10. For (B-C) the dashed red lines delimits the 1% most extreme differences between replicates which were removed in the ART modelling approach. (D) GFP synthesis rate compared to strain genotype (n = 3). The data is ordered according to decreasing mean GFP synthesis rate. Data points included in the TeselaGen EVOLVE modeling approach are shown in green, whereas data points in red or black were excluded. Red markers indicate outliers whereas black markers indicates strains for which only one replicate is left after outlier removal.

**Figure S6. Construction of an easy-curable plasmid using counter selection**. Two dosage sensitive genes (*ACT1* & *CDC14*) were expressed under the control of the galactose-inducible *GAL1* promoter and cloned into USER vector pRS413-mKate2 (pCfB2866, Zhang et al., 2016). To test the efficiency of counter selection, yeast strain with a plasmid containing one of the counter selection cassettes (pRS413-HIS3 P_*GAL1*_-*ACT1*-T_*IDP1*_ or P_*GAL1*_-*CDC14*-T_*ADH1*_) was grown in both non-induction (synthetic complete + glucose) and induction (synthetic complete + galactose) media for 18 hrs. A diluted aliquot of culture was spread onto both YPD (without selection for the *HIS3* selectable marker) and SC-HIS (with selection for the *HIS3* selectable marker) drop out agar plates. Only cultures without growth on SC-HIS selective media were used for further studies.

**Table S1**. Primers used in study. Sequence features of interest are separated by a space.

**Table S2**. Plasmids constructed and used in study.

**Table S3**. Yeast strains engineered and used in study.

**Table S4**. Related to Figure 1. Gene scores of all 192 genome-scale modelled (FBA) genes with significant changes in flux towards tryptophan production under glucose and ethanol conditions. A score higher than one means the gene is an up-regulation candidate, a score between zero and one means the gene is a down-regulation candidate, a score equal to zero means the gene is a knockout candidate, and a blank score means the gene is associated to reactions that do not change significantly in flux as tryptophan production increases under that particular condition. The four out of five gene targets identified by FBA and selected for this study are marked in bold.

**Table S5**. Related to Figure 1. FBA results for all pathways in metabolism, including the number of gene targets predicted in each pathway, the total size of each pathway, the fraction of genes in each pathway that are gene targets, and the significance of that representation in each pathway compared to the rest of metabolism (“Whole metabolism”), indicated by a P-value computed with a Fisher’s exact test.

**Table S6**. Related to Figure 1. The 30 selected native yeast promoters, and their position in the combinatorial cluster.

**Table S7**. Related to Figure 3D. Promoter combinations of library control strains. The numbers in each row refer to promoter numbers as shown in Table S5. Design no. 1 contains the promoters that are native to the genes at the five positions.

**Table S8**. Related to Figure 1 and 4C. Top-30 promoter combinations as recommended by ART. Size of color bars indicate promoter expression strength (see Figure 1), and column “dgfp/dt” shows predicted GFP synthesis rate.

**Table S9**. Related to Figure 1 and 4C. Top-30 promoter combinations as recommended by TeselaGen EVOLVE. Size of color bars indicate promoter expression strength (see Figure 1), and column “dgfp/dt” shows predicted GFP synthesis rate.

