## Supplementary information for "Predictive engineering and optimization of tryptophan metabolism in yeast through a combination of mechanistic and machine learning models"

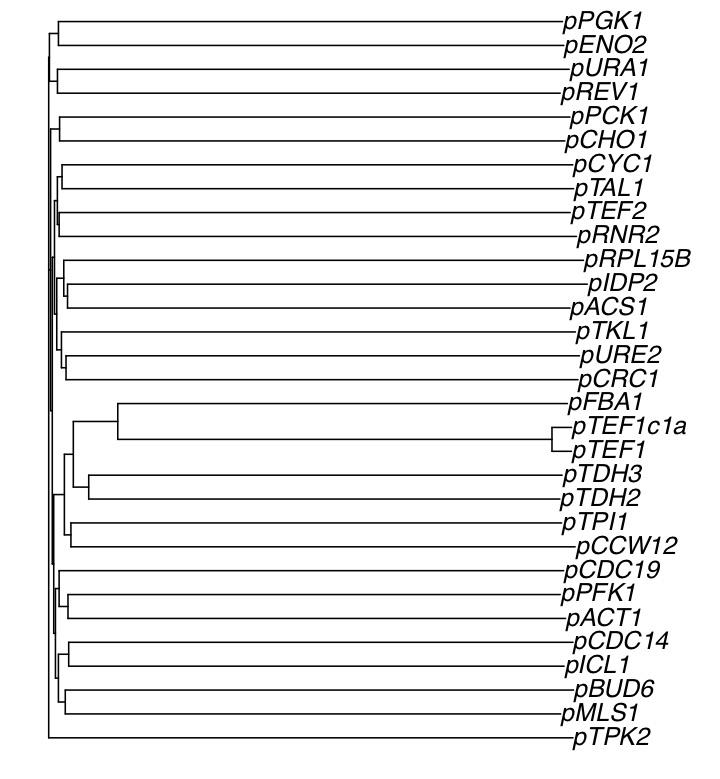

**Figure S1. Related to Figure 1. Dendrogram of the sequence diversity of 30 selected native yeast promoters.** Sequence pTEF1c1a with a single nucleotide change from pTEF1 has been added as a reference. The dendrogram was constructed using the neighbor-joining method (Saitou and Nei, 1987; Studier and Keppler, 1988).

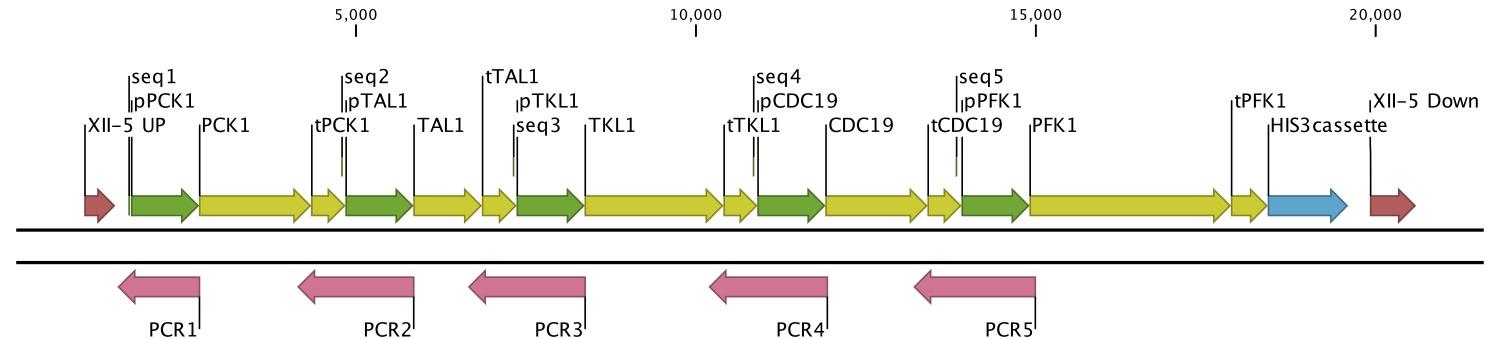

**Figure S2. Related to Figure 1. Genotyping strategy.** Schematic outline of the genotyping strategy to assess correct *in vivo* junction-junction assemblies of 11 parts, and the integration at EasyClone site XII-5 (Jensen et al., 2014). Marked in red are chromosomal regions of EasyClone site XII-5, whereas green marks the promoters, and yellow the coding sequences and terminators. Marked in blue is the selectable *HIS3* expression cassette, while genotyping PCRs are marked in light red. Primers used for sequencing of the 5 PCR reactions are marked seq1-seq5.

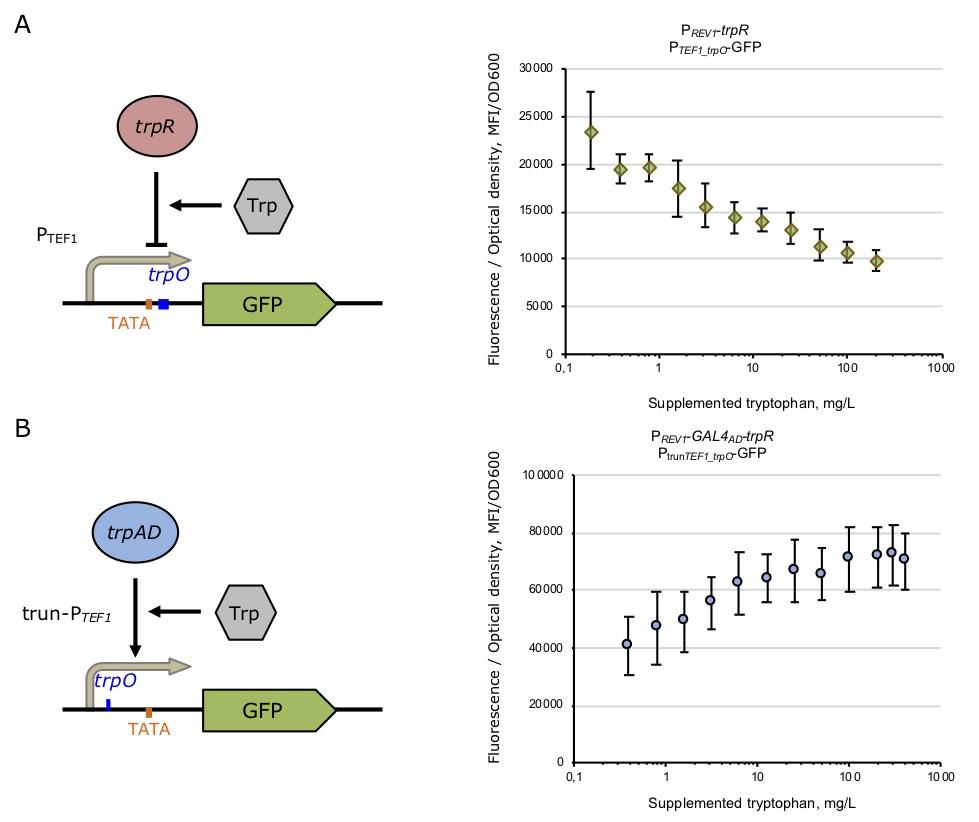

**Figure S3. Related to Figure 3. Biosensor development and characterization.** Overnight cultures of the strain containing sensor and reporter was used to inoculate fresh media supplemented with various concentrations of tryptophan and grown for 6 hours (early-mid exponential phase). Optical density (measured as absorbance at 600 nm) was used to normalize the green fluorescence (excitation/emission at 485/515 nm). (A) *E. coli* *trpR* was directly expressed in a yeast strain harboring the yEGFP reporter under the control of *TEF1* promoter containing *trpO* sequence inserted downstream of the TATA-like element. (B) The *trpR* gene was fused to the C-terminus of the activator domain of GAL4 (GAL4_ad_) with a GSGSGS linker, turning this transcriptional repressor into an activator (trpAD). Accordingly, the *trpO* sequence was placed upstream of a truncated *TEF1* promoter (lacking region with multiple Rap1-binding sites).

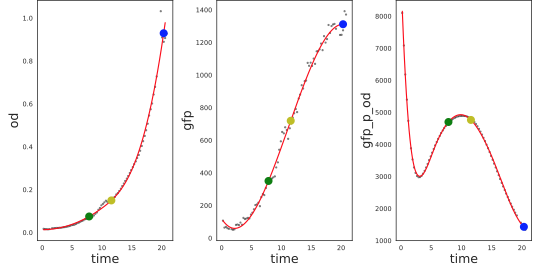

**Figure S4. Related to Figure 3E-F. Parameter estimation from time series data.** (A) Representative growth curve of *S. cerevisiae* in microtiter plates. *S. cerevisiae* was grown in yeast synthetic drop-out media in 96-well microtiter plates, and cell density measured at 600 nm (OD_600_) over 20 hrs. (B) Representative tryptophan biosensor output measured as fluorescence (GFP) in *S. cerevisiae* cells (n = 1). *S. cerevisiae* was grown in yeast synthetic drop-out media in 96-well microtiter plates, and GFP measured at 485 nm (OD_485_) over 20 hrs. (C) Tryptophan biosensor output normalized by absorbance at 600 nm (OD_600_) over 20 hrs. For (A-C) the red line shows model fitting using a univariate spline. All plots represent a single replicate measurement (n = 1). The green, yellow and blue markers indicate OD_600_ = 0.075, OD_600_ = 0.15, and maximum rate of OD_600_ increase, respectively.

When calculating GFP synthesis rates (increase in GFP/time) we normalized our measurements with the number of cells (GFP/OD_600_/time), because it is not possible to inoculate the medium with exactly the same number of cells. In order to calculate normalized rates (GFP/OD_600_) we measured both OD_600_ and GFP over time for all >500 strains. We only calculated rates in the period when GFP/OD_600_ was fairly constant and high (Figure 3E, Figure S4). Here, we observed that this was the case in the early part of the exponential phase, i.e. not in the entire exponential growth phase. From this, we observed that increase in GFP/time declined before OD_600_/time. This is considered to be due GFP maturation being more sensitive to oxygen than to cell growth. Picking the correct period for calculating rates was necessary to make sure that we got the actual strain characteristics, and not biases due to the specific laboratory setup (e.g. that the cells begin to shade one another at high OD_600_ and thereby limit detection of GFP, or due to feedback degradation of GFP). By ensuring this we achieved high reproducibility, and thus a higher signal to noise ratio (Figure 3F).

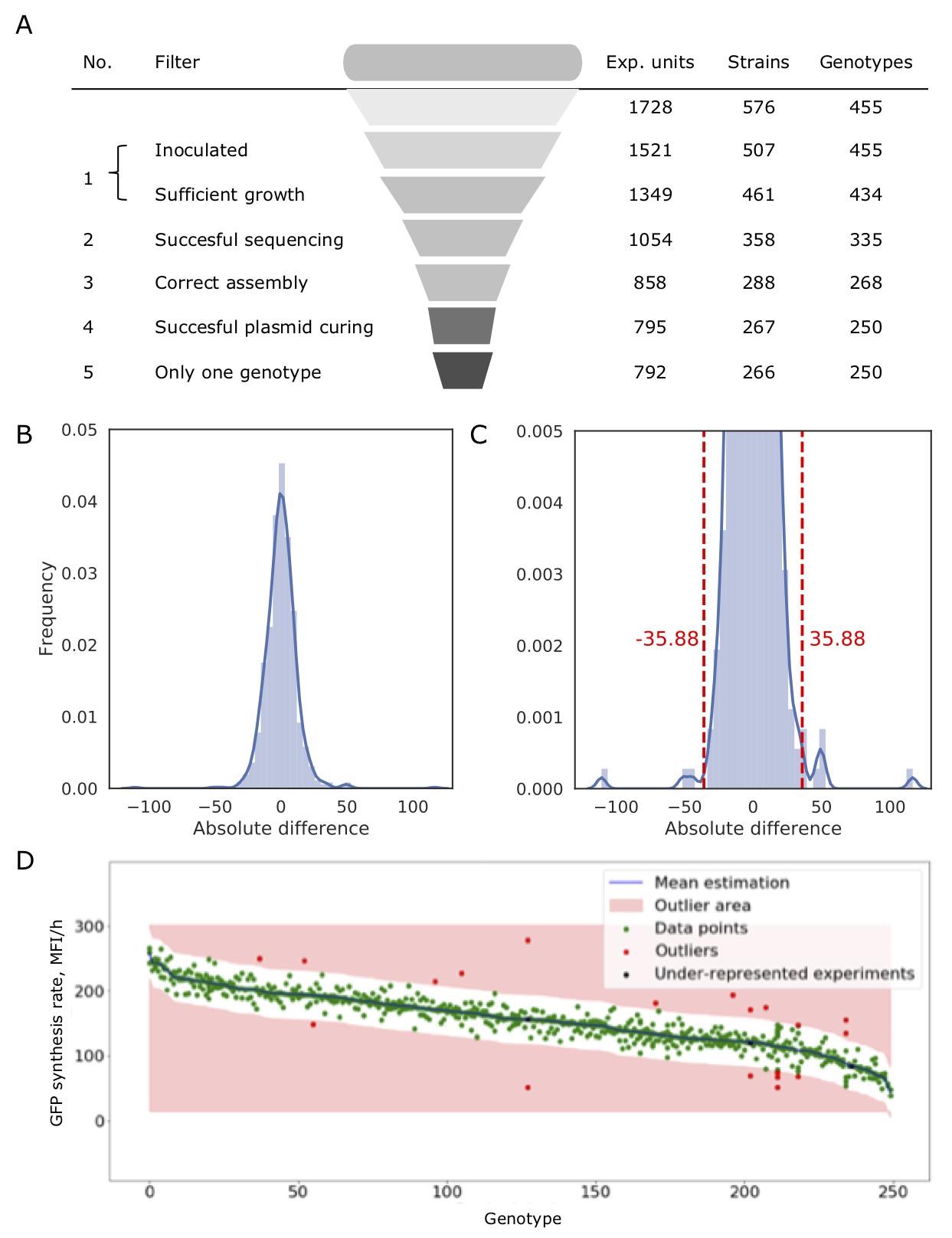

**Figure S5. Related to Figures 3-4. Data filtering and outlier removal.** (A) Schematic illustration of the various filtering steps applied for data quality control. The six steps used for filtering are indicated by number to the left, and listed to the right are the numbers of unique genotypes as inferred from sequencing, the number of strains, and the number of experimental units (Exp. units, n = 3). (B) The distribution of absolute differences between replicate measurements (n = 3) of strain GFP synthesis rate. (C) Same as in (B), but with y-axis expanded by a factor 10. For (B-C) the dashed red lines delimits the 1% most extreme differences between replicates which were removed in the ART modelling approach. (D) GFP synthesis rate compared to strain genotype (n = 3). The data is ordered according to decreasing mean GFP synthesis rate. Data points included in the TeselaGen EVOLVE modeling approach are shown in green, whereas data points in red or black were excluded. Red markers indicate outliers whereas black markers indicates strains for which only one replicate is left after outlier removal.

**Figure S6. Construction of an easy-curable plasmid using counter selection.** Two dosage sensitive genes (*ACT1* & *CDC14*) were expressed under the control of the galactose-inducible *GAL1* promoter and cloned into USER vector pRS413-mKate2 (pCfB2866, [Zhang et al., ACS Synth Biol](https://pubs.acs.org/doi/abs/10.1021/acssynbio.6b00135)). To test the efficiency of counter selection, yeast strain with a plasmid containing one of the counter selection cassettes (pRS413-HIS3 P*_GAL1_-ACT1*-T*_IDP1_* or P*_GAL1_-CDC14*-T*_ADH1_*) was grown in both non-induction (synthetic complete + glucose) and induction (synthetic complete + galactose) media for 18 hrs. A diluted aliquot of culture was spread onto both YPD (without selection for the *HIS3* selectable marker) and SC-HIS (with selection for the *HIS3* selectable marker) drop out agar plates. Only cultures without growth on SC-HIS selective media were used for further studies.
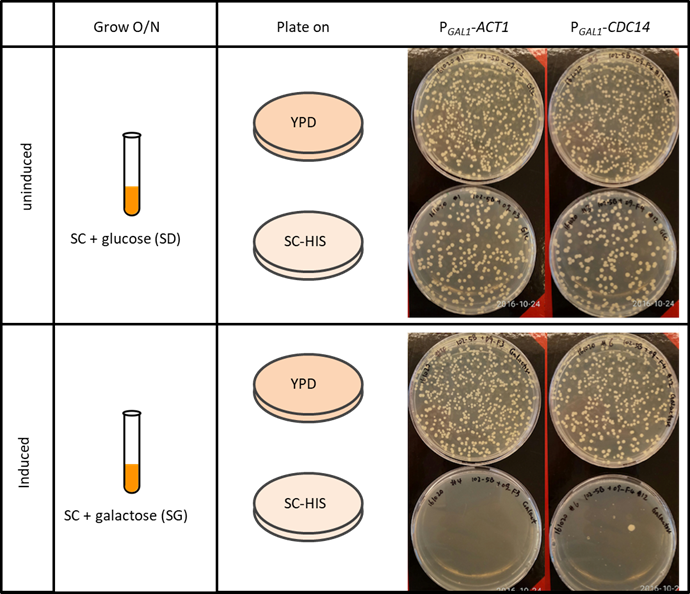

**Table S1**. Primers used in study. Sequence features of interest are separated by a space.

| **Name** | **Sequence (5' - 3')** | **Use** |
| --- | --- | --- |
| **Library construction** | | |
| tADH1-pPCK1_Fw | CTTATTTAGAAGTGTCAACAACGTATCTAC  CACATGTCGACGAGTTT | Forward pPCK1, overhang to EasyClone site XII-5 UP |
| PCK1_N-pPCK1_Rv | TACTGTAGCATTCATTTTAGAAGGGGACAT  GTTGTTATTTTATTATGGAATAATTAGT | Reverse pPCK1, overhang to PCK1 N-terminal |
| tADH1-pTPI1_Fw | CTTATTTAGAAGTGTCAACAACGTATCTAC  AAGGATGAGCCAAGAATAA | Forward pTPI1, overhang to EasyClone site XII-5 UP |
| PCK1_N-pTPI1_Rv | TACTGTAGCATTCATTTTAGAAGGGGACAT  TTTTAGTTTATGTATGTGTTTTTTG | Reverse pTPI1, overhang to PCK1 N-terminal |
| tADH1-pICL1_Fw | CTTATTTAGAAGTGTCAACAACGTATCTAC  TTGGAAATGTAAAGGATAAT G | Forward pICL1, overhang to EasyClone site XII-5 UP |
| PCK1_N-pICL1_Rv | TACTGTAGCATTCATTTTAGAAGGGGACAT  TTTTCGTTGACTTTTTGTTAT | Reverse pICL1, overhang to PCK1 N-terminal |
| tADH1-pRNR2_Fw | CTTATTTAGAAGTGTCAACAACGTATCTAC  TTCTTTATCTTTTTTTCCCTT | Forward pRNR2, overhang to EasyClone site XII-5 UP |
| PCK1_N-pRNR2_Rv | TACTGTAGCATTCATTTTAGAAGGGGACAT  GGTAATTGGACAAATAAATACG | Reverse pRNR2, overhang to PCK1 N-terminal |
| tADH1-pACT1_Fw | CTTATTTAGAAGTGTCAACAACGTATCTAC  ACCCATATAATATAATAACTAAATAAGTAA | Forward pACT1, overhang to EasyClone site XII-5 UP |
| PCK1_N-pACT1_Rv | TACTGTAGCATTCATTTTAGAAGGGGACAT ACCAGAACCGTTATCAATAAC | Reverse pACT1, overhang to PCK1 N-terminal |
| tADH1-pTHD3_Fw | CTTATTTAGAAGTGTCAACAACGTATCTAC  CTATTTTCGAGGACCTTGT | Forward pTDH3, overhang to EasyClone site XII-5 UP |
| PCK1_N-pTHD3_Rv | TACTGTAGCATTCATTTTAGAAGGGGACAT  TTTGTTTGTTTATGTGTGTTTAT | Reverse pTDH3, overhang to PCK1 N-terminal |
| tPCK1-pTAL1_Fw | ACTTGTATCGAATCACTTTACCGTTCTTTA  TCTATACCCATTGATCGG | Forward pTAL1, overhang to tPCK1 |
| TAL1_N-pTAL1_Rv | CTTTTGTTTCTTTTGAGCTGGTTCAGACAT  TATGTACACGTATATGTGACGA | Reverse pTAL1, overhang to TAL1 N-terminal |
| tPCK1-pENO2_Fw | ACTTGTATCGAATCACTTTACCGTTCTTTA  AATCCTACTCTTGCCGTT | Forward pENO2, overhang to tPCK1 |
| TAL1_N-pENO2_Rv | CTTTTGTTTCTTTTGAGCTGGTTCAGACAT  TATTATTGTATGTTATAGTATTAGTTGCTT | Reverse pENO2, overhang to TAL1 N-terminal |
| tPCK1-pACS1_Fw | ACTTGTATCGAATCACTTTACCGTTCTTTA  TGGTCTGCAAATGCTTT | Forward pACS1, overhang to tPCK1 |
| TAL1_N-pACS1_Rv | CTTTTGTTTCTTTTGAGCTGGTTCAGACAT  AACACAGTGGGGCAAT | Reverse pACS1, overhang to TAL1 N-terminal |
| tPCK1-pREV1_Fw | ACTTGTATCGAATCACTTTACCGTTCTTTA  TTCTTAGGCACAACAATATTTATA | Forward pREV1, overhang to tPCK1 |
| TAL1_N-pREV1_Rv | CTTTTGTTTCTTTTGAGCTGGTTCAGACAT  CGCTGGATATGCCTAGA | Reverse pREV1, overhang to TAL1 N-terminal |
| tPCK1-pCHO1_Fw | ACTTGTATCGAATCACTTTACCGTTCTTTA AAAAAGCGAATCATGTAGAT | Forward pCHO1, overhang to tPCK1 |
| TAL1_N-pCHO1_Rv | CTTTTGTTTCTTTTGAGCTGGTTCAGACAT AACGTCTGTGTCCGTG | Reverse pCHO1, overhang to TAL1 N-terminal |
| tPCK1-pCYC1_Fw | ACTTGTATCGAATCACTTTACCGTTCTTTA CAGCATTTTCAAAGGTGT | Forward pCYC1, overhang to tPCK1 |
| TAL1_N-pCYC1_Rv | CTTTTGTTTCTTTTGAGCTGGTTCAGACAT TATTAATTTAGTGTGTGTATTTGTGT | Reverse pCYC1, overhang to TAL1 N-terminal |
| tTAL1-pTKL1_Fw | CCATTTTGACCTGAATCAAACGAATGAATG ATACTGGACTTGGAAATTCC | Forward pTKL1, overhang to tTAL1 |
| TKL1_N-pTKL1_Rv | TAGCTTATCAATGTCAGTGAATTGAGTCAT TTTGTTTGCTAAAGAGGTAACT | Reverse pTKL1, overhang to TKL1 N-terminal |
| tTAL1-pPGK1_Fw | CCATTTTGACCTGAATCAAACGAATGAATG AGACGCGAATTTTTCGA | Forward pPGK1, overhang to tTAL1 |
| TKL1_N-pPGK1_Rv | TAGCTTATCAATGTCAGTGAATTGAGTCAT TGTTTTATATTTGTTGTAAAAAGTAGATAA | Reverse pPGK1, overhang to TKL1 N-terminal |
| tTAL1-pMLS1_Fw | CCATTTTGACCTGAATCAAACGAATGAATG TTTAATCTTTAGGGAGGGTAA | Forward pMLS1, overhang to tTAL1 |
| TKL1_N-pMLS1_Rv | TAGCTTATCAATGTCAGTGAATTGAGTCAT TTTCTTAATTCTTTTATGTGCTT | Reverse pMLS1, overhang to TKL1 N-terminal |
| tTAL1-pBUD6_Fw | CCATTTTGACCTGAATCAAACGAATGAATG CTTTTGAAGACTGCTGCT | Forward pBUD6, overhang to tTAL1 |
| TKL1_N-pBUD6_Rv | TAGCTTATCAATGTCAGTGAATTGAGTCAT CTAATTTTAAAATAATACGAGGATTAC | Reverse pBUD6, overhang to TKL1 N-terminal |
| tTAL1-pURE2_Fw | CCATTTTGACCTGAATCAAACGAATGAATG CAAGCTGAACTCGCTGA | Forward pURE2, overhang to tTAL1 |
| TKL1_N-pURE2_Rv | TAGCTTATCAATGTCAGTGAATTGAGTCAT TTGGTGTAGAACTTAATTTGC | Reverse pURE2, overhang to TKL1 N-terminal |
| tTAL1-pCCW12_Fw | CCATTTTGACCTGAATCAAACGAATGAATG AAAGAAACTTAATACGTTATGCC | Forward pCCW12, overhang to tTAL1 |
| TKL1_N-pCCW12_Rv | TAGCTTATCAATGTCAGTGAATTGAGTCAT TATTGATATAGTGTTTAAGCGAA | Reverse pCCW12, overhang to TKL1 N-terminal |
| tTKL1-pCDC19_Fw | AGTCGAAAAGGCTAATCTAGAAAATCGATT ACTTGAGATGTGTGTCAATG | Forward pCDC19, overhang to tTKL1 |
| CDC19_N-pCDC19_Rv | TAATGAGGTCAATCTTTCTAATCTAGACAT TGTGATGATGTTTTATTTGTTT | Reverse pCDC19, overhang to CDC19 N-terminal |
| tTKL1-pURA1_Fw | AGTCGAAAAGGCTAATCTAGAAAATCGATT GTTGTATTAATTTTCTCGAAGG | Forward pURA1, overhang to tTKL1 |
| CDC19_N-pURA1_Rv | TAATGAGGTCAATCTTTCTAATCTAGACAT GTTTGGTACGGAAGTTCA | Reverse pURA1, overhang to CDC19 N-terminal |
| tTKL1-pCRC1_Fw | AGTCGAAAAGGCTAATCTAGAAAATCGATT TAGTTGATTTATTTCCCTGC | Forward pCRC1, overhang to tTKL1 |
| CDC19_N-pCRC1_Rv | TAATGAGGTCAATCTTTCTAATCTAGACAT TACTGACACGATGACGTTT | Reverse pCRC1, overhang to CDC19 N-terminal |
| tTKL1-pCDC14_Fw | AGTCGAAAAGGCTAATCTAGAAAATCGATT GTTGTGTATTTCGTACCTATGTAT | Forward pCDC14, overhang to tTKL1 |
| CDC19_N-pCDC14_Rv | TAATGAGGTCAATCTTTCTAATCTAGACAT TTATAAGCGTACTTTGTAGTCC | Reverse pCDC14, overhang to CDC19 N-terminal |
| tTKL1-pTEF2_Fw | AGTCGAAAAGGCTAATCTAGAAAATCGATT GTAGGTGTTCCTTGAGCTAC | Forward pTEF2, overhang to tTKL1 |
| CDC19_N-pTEF2_Rv | TAATGAGGTCAATCTTTCTAATCTAGACAT GTTTAGTTAATTATAGTTCGTTGACC | Reverse pTEF2, overhang to CDC19 N-terminal |
| tTKL1-pFBA1_Fw | AGTCGAAAAGGCTAATCTAGAAAATCGATT ACTGGTAGAGAGCGACTTT | Forward pFBA1, overhang to tTKL1 |
| CDC19_N-pFBA1_Rv | TAATGAGGTCAATCTTTCTAATCTAGACAT TTTGAATATGTATTACTTGGTTATG | Reverse pFBA1, overhang to CDC19 N-terminal |
| tCDC19-pPFK1_Fw | ACGCGGGCAGATTCAATTAGTGTCCTAAAT ACCTCATCTATAATTTTTACCCT | Forward pPFK1, overhang to tCDC19 |
| PFK1_N-pPFK1_Rv | AACACCGTAGCATGAATCTTGAGATTGCAT CTTTGATATGATTTTGTTTCAG | Reverse pPFK1, overhang to PFK1 N-terminal |
| tCDC19-pTDH2_Fw | ACGCGGGCAGATTCAATTAGTGTCCTAAAT CTAGATCAGAGGGTGGTAAAT | Forward pTDH2, overhang to tCDC19 |
| PFK1_N-pTDH2_Rv | AACACCGTAGCATGAATCTTGAGATTGCAT TTTGTTTTGTTTGTTTGTGT | Reverse pTDH2, overhang to PFK1 N-terminal |
| tCDC19-pIDP2_Fw | ACGCGGGCAGATTCAATTAGTGTCCTAAAT AATAGTCTTACACCAATGAGC | Forward pIDP2, overhang to tCDC19 |
| PFK1_N-pIDP2_Rv | AACACCGTAGCATGAATCTTGAGATTGCAT TACGATTTTATATATATACGTACGTTAC | Reverse pIDP2, overhang to PFK1 N-terminal |
| tCDC19-pTPK2_Fw | ACGCGGGCAGATTCAATTAGTGTCCTAAAT CAACAAGTCTGAAACTTTCA | Forward pTPK2, overhang to tCDC19 |
| PFK1_N-pTPK2_Rv | AACACCGTAGCATGAATCTTGAGATTGCAT ACCGACAATTTTCAACAG | Reverse pTPK2, overhang to PFK1 N-terminal |
| tCDC19-pRPL15B_Fw | ACGCGGGCAGATTCAATTAGTGTCCTAAAT GTACTGCTGGCCATTTTTAT | Forward pRPL15B, overhang to tCDC19 |
| PFK1_N-pRPL15B_Rv | AACACCGTAGCATGAATCTTGAGATTGCAT TGCTTGTGTGGTAGGTAATT | Reverse pRPL15B, overhang to PFK1 N-terminal |
| tCDC19-pTEF1_Fw | ACGCGGGCAGATTCAATTAGTGTCCTAAAT CTTCATCGGTATCTTCGC | Forward pTEF1, overhang to tCDC19 |
| PFK1_N-pTEF1_Rv | AACACCGTAGCATGAATCTTGAGATTGCAT TTTGTAATTAAAACTTAGATTAGATTG | Reverse pTEF1, overhang to PFK1 N-terminal |
| PCK1_N_Fw | ATGTCCCCTTCTAAAATGA | Forward PCK1 |
| tPCK1_Rv-1 | TAAAGAACGGTAAAGTGATTC | Reverse PCK1 |
| TAL1_N_Fw | ATGTCTGAACCAGCTCAA | Forward TAL1 |
| tTAL1_Rv-1 | CATTCATTCGTTTGATTCA | Reverse TAL1 |
| TKL1_N_Fw | ATGACTCAATTCACTGACATT | Forward TKL1 |
| tTKL1_Rv-1 | AATCGATTTTCTAGATTAGCC | Reverse TKL1 |
| CDC19_N_Fw | ATGTCTAGATTAGAAAGATTGACC | Forward CDC19 |
| tCDC19_Rv-1 | ATTTAGGACACTAATTGAATCTG | Reverse CDC19 |
| PFK1_N_Fw | ATGCAATCTCAAGATTCATG | Forward PFK1 |
| tPFK1_Rv-1 | CACTAGTTTCCATTTTTCCA | Reverse PFK1 |
| EC_UP_Fw | AAAGTATAGGAACTTCTGAAGTGG | Forward XII-5 UP |
| tADH1_Rv | GTAGATACGTTGTTGACACTTCTAAATA | Reverse tADH1 |
| tCYC1_Fw | ATCCGCTCTAACCGAAAAG | Forward tCYC1 |
| EC_DW_Rv | AACTTCACTTCATTTTATTTAAATTTGC | Reverse XII-5 DW |
| tPFK1_pHIS3_Fw | GTTTCTTTTTATCTTTCCGCTGGAAAAATGGAAACTAGTG CGTTTTAAGAGCTTGGTGAG | Forward pHIS3, overhang to tPFK1 |
| tCYC1_tHIS3_Rv | CTAACTCCTTCCTTTTCGGTTAGAGCGGAT NNNNNNNNN ATAGATCCGTCGAGTTCAAGA | Reverse tHIS3, overhang to tCYC1 |
| **Sequencing for construction validation** | | |
| XII-5-up-out-sq | CCACCGAAGTTGATTTGCTT | Forward upstream sequence integrated at EasyClone site XII-5 |
| TADH1_towards out | GTTGACACTTCTAAATAAGCGAATTTC | Reverse beginning of integrated sequencing at EasyClone site XII-5 |
| DW_towards out | CCTGCAGGACTAGTGCTGAG | Forward end of integrated sequence at EasyClone site XII-5 |
| XII-5-down-out-sq | GTGGGAGTAAGGGATCCTGT | Reverse downstream sequence integrated at EasyClone site XII-5 |
| USER_XhoI_Fw | ACTCTCGAG AGCGACCTCATGCTATACC | Forward tADH1, designed to test junctions around promoter at position 1 |
| PCK1_N_Rv | ATTCATTTTAGAAGGGGACAT | Reverse N-terminal of PCK1, designed to test junctions around promoter at position 1 |
| PCK1_C_Fw | CGATTTTCAATCTTCAAGTAC | Forward C-terminal PCK1, designed to test junctions around promoter at position 2 |
| TAL1_N_Rv | TTTGAGCTGGTTCAGACAT | Reverse N-terminal of TAL1, designed to test junctions around promoter at position 2 |
| TAL1_C_Fw | CTTTCCCAAGAGTTTTGG | Forward C-terminal TAL1, designed to test junctions around promoter at position 3 |
| TKL1_N_Rv | CAATGTCAGTGAATTGAGTCAT | Reverse N-terminal of TKL1, designed to test junctions around promoter at position 3 |
| TKL1_C_Fw-2 | TCCAATCATGTCTGTTGAA | Forward C-terminal TKL1, designed to test junctions around promoter at position 4 |
| CDC19_N_Rv-2 | CAGCAACAACGTTTAATGA | Reverse N-terminal of CDC19, designed to test junctions around promoter at position 4 |
| CDC19_C_Fw | ACTTGTACAGAGGTGTCTTCC | Forward C-terminal CDC19, designed to test junctions around promoter at position 5 |
| PFK1_N_Rv | CTAGAGTGTGATAAAAGTGAATG | Reverse N-terminal of PFK1, designed to test junctions around promoter at position 5 |
| ADH1_test_fw | GAAATTCGCTTATTTAGAAGTGTC | Forward tADH1, designed to identify promoter at position 1 |
| seq_junction_2 | GGTTACCGGAATGATTCACCG | Forward tPCK1, designed to identify promoter at position 2 |
| seq_junction_3 | CGATGCTGTAAACGTCCCTG | Forward tTAL1, designed to identify promoter at position 3 |
| seq_junction_4 | GATCACCAATGGCGGAAGC | Forward tTKL1, designed to identify promoter at position 4 |
| seq_junction_5 | GTTCAGCTTCTGGCCTTCG | Forward tCDC19, designed to identify promoter at position 5 |

**Table S2.** Plasmids constructed and used in study.

| **Name** | **Description** | **Reference** |
| --- | --- | --- |
| **Tryptophan biosensor development** | | |
| pCfB4107 | CEN6/ARS4 pRS413U-*HIS3*, P*_TEF2_trpO_*-*yEGFP*-T*_ADH1_* | This study |
| pCfB4108 | CEN6/ARS4 pRS416U-*HIS3*, P*_REV1_*-*trpR*-T*_ADH1_* | This study |
| pCfB4743 | CEN6/ARS4 pRS416U-*URA3*, P*_REV1_*-*GAL4*_ad__*trpR*-T*_ADH1_* | This study |
| pCfB4747 | CEN6/ARS4 pRS416U-*URA3*, P*_REV1_*-*GAL4*_ad_-T*_ADH1_* | This study |
| pCfB4750 | CEN6/ARS4 pRS413U-*HIS3*, P*_TEF2_*-*mKate2*-T*_IDP1_*, P*_trunTEF1_trpO_*-yEGFP-T*_ADH1_* | This study |
| pCfB5397 | CEN6/ARS4, pRS413U-*HIS3*, P*_GAL1core_3xtrpO_*-*yEGFP*-T*_ADH1_*, P*_TEF1_trpO_*-*mKate2*-T*_CYC1_* | This study |
| pCfB5399 | CEN6/ARS4, pRS413U-*HIS3*, P*_GAL1core_6xtrpO_*-*yEGFP*-T*_ADH1_*, P*_TEF1_trpO_*-*mKate2*-T*_CYC1_* | This study |
| **Platform and library strain construction** | | |
| pCfB176 | CEN6/ARS4, pRS414-*TRP1*, P*_TEF1_*-SpCas9-T*_CYC1_* | DiCarlo et al., 2013 |
| pCfB4672 | CEN6/ARS4, pRS413U-*HIS3*, P*_TEF2_*-*mKate2*-T*_IDP1_*, P*_GAL1_*-*ACT1*-T*_ADH1_* | This study |
| pCfB4673 | CEN6/ARS4, pRS413U-*HIS3*, P*_TEF2_*-*mKate2*-T*_IDP1_*, P*_GAL1_*-*CDC14*-T*_ADH1_* | This study |
| pCfB9303 | CEN6/ARS4, pRS415U-*LEU2*, P*_GAL1_*-*ACT1*-T*_IDP1_* | This study |
| pCfB9307 | CEN6/ARS4, pRS415U*-LEU2*, P*_GAL1_*-*ACT1*-T*_IDP1_*, *TKL1-TAL1-PFK1-CDC19* (native expression cassettes) | This study |
| pCfB6842 | 2 *µ*, pESC-*LEU2,* P*_SNR52_*-*ARO4*_gRNA-T*_SUP4_* | This study |
| pCfB6843 | 2 *µ*, pESC-*LEU2,* P*_SNR52_*-*ARO4*_gRNA-T*_SUP4_*, P*_SNR52_*-*TRP2*_gRNA_1-T*_SUP4_*, P*_SNR52_*-*TRP2*_gRNA_2-T*_SUP4_* | This study |
| pCfB6844 | 2 *µ*, pESC-*LEU2,* P*_SNR52_*-*TRP2*_gRNA_1-T*_SUP4_*, P*_SNR52_*-*TRP2*_gRNA_2-T*_SUP4_* | This study |
| pCfB6903 | 2 *µ*, pESC-*LEU2,* P*_SNR52_*-XI-2_gRNA-T*_SUP4_* | This study |
| pCfB6904 | 2 *µ*, pESC-*LEU2,* P*_SNR52_*-XI-3_gRNA-T*_SUP4_* | This study |
| pCfB6909 | 2 *µ*, pESC-*LEU2*-P*_SNR52_*-XII-5_gRNA-T*_SUP4_* | This study |
| pCfB6916 | 2 *µ*, pESC-*URA3,* P*_SNR52_*-XI-5_gRNA-T*_SUP4_* | This study |
| pCfB6895 | 2 µ, pESC-*URA3*, P*_SNR52_*-*PCK1*_gRNA-T*_SUP4,_* P*_SNR52_*-*TAL1*_gRNA_1-T*_SUP4,_* P*_SNR52_*-*TAL1*_gRNA_2-T*_SUP4,_* P*_SNR52_*-*TKL1*_gRNA-T*_SUP4_* | This study |
| pCfB9306 | 2 µ, pESC-*URA3*, P*_SNR52_*-*pPFK1*_gRNA_1-T*_SUP4,_* P*_SNR52_*-*pPFK1*_gRNA_2-T*_SUP4,_* P*_SNR52_*-p*CDC19*_gRNA_1-T*_SUP4,_* P*_SNR52_*-p*CDC19*_gRNA_2-T*_SUP4_* | This study |

**Table S3.** Yeast strains engineered and used in study.

| **Name** | **Genotype** | **Reference** |
| --- | --- | --- |
| CEN.PK113-11C | *MAT***a** *his3*∆1, *LEU2*, *ura3*-52, *TRP1* *MAL2*-8c *SUC2* | EUROSCARF |
| CEN.PK2-1C | *MAT***a** *his3*∆1, *leu2*-3_112, *ura3*-52, *trp1*-289, *MAL2*-8c *SUC2* | EUROSCARF |
| TrpA-1 | *MAT***a** P*_GAL1core_6xtrpO_*-*yEGFP*-T*_ADH1_*, P*_TEF1_trpO_*-*mKate2*-T*_CYC1_*, pCfB176 | this study |
| TrpA-2 | *MAT***a** P*_GAL1core_6xtrpO_*-*yEGFP*-T*_ADH1_*, P*_TEF1_trpO_*-mKate2-T*_CYC1_*, *ARO4^wt^*::*ARO4*^K229L^, pCfB176 | this study |
| TrpA-3 | *MAT***a** P*_GAL1core_6xtrpO_*-*yEGFP*-T*_ADH1_*, P*_TEF1_trpO_*-mKate2-T*_CYC1_, TRP2^wt^*::*TRP2*^S65R, S76L^, pCfB176 | this study |
| TrpA-4 | *MAT***a** P*_GAL1core_6xtrpO_*-*yEGFP*-T*_ADH1_*, P*_TEF1_trpO_*-mKate2-T*_CYC1_*, *ARO4^wt^*::*ARO4*^K229L^, *TRP2^wt^*::*TRP2*^S65R, S76L^, pCfB176 | this study |
| TrpNA-W | *MAT***a** *tkl1*∆ *tal1*∆ *pck1*∆, P*_PFK1_*::P*_REV1_*-*PFK1*,  P*_CDC19_*::P*_RNR2_*-*CDC19*, P*_PFK1_*-*GAL4*_ad_-*trpR*-T*_ADH1_*,  P*_GAL1core_3xtrpO_*-*yEGFP*-T*_ADH1_*, P*_TEF1_trpO_*-*mKate2*-T*_CYC1_*, P*_PGK1_*-*ARO4^K229L^*-T*_ADH1_*,  P*_TEF1_*-*TRP2*^S65R, S76L^-T*_CYC1_*, pCfB176, pCfB9307 | this study |

**Table S4**. Related to Figure 1. Gene scores of all 192 genome-scale modelled (FBA) genes with significant changes in flux towards tryptophan production under glucose and ethanol conditions. A score higher than one means the gene is an up-regulation candidate, a score between zero and one means the gene is a down-regulation candidate, a score equal to zero means the gene is a knockout candidate, and a blank score means the gene is associated to reactions that do not change significantly in flux as tryptophan production increases under that particular condition. The four out of five gene targets identified by FBA and selected for this study are marked in bold.

| **Gene name** | **Glucose** | **Ethanol** |  | **Gene name** | **Glucose** | **Ethanol** |  | **Gene name** | **Glucose** | **Ethanol** |
| --- | --- | --- | --- | --- | --- | --- | --- | --- | --- | --- |
| YLR438W | 1000 | 0 |  | YER070W | 0,51899 | 0,433024 |  | YML008C |  | 17,67071 |
| YBR249C | 636,7273 |  |  | YGR180C | 0,51899 | 0,433024 |  | YGL055W |  | 6,313433 |
| YKL120W | 636,7273 |  |  | YIL066C | 0,51899 | 0,433024 |  | YKL182W |  | 4,098839 |
| YHR208W | 500,275 | 0,129706 |  | YJL026W | 0,51899 | 0,433024 |  | YPL231W |  | 4,098839 |
| YBR068C | 448,3277 | 353,2292 |  | YBR218C | 0,5093 | 636,7273 |  | YDR353W |  | 2,608208 |
| YBR069C | 448,3277 | 353,2292 |  | YGL062W | 0,5093 | 636,7273 |  | YLR058C |  | 2,443903 |
| YDR046C | 448,3277 | 353,2292 |  | YEL039C | 0,49376 |  |  | **YKR097W (PCK1)** |  | **1,576565** |
| YKR039W | 448,3277 | 353,2292 |  | YJR048W | 0,49376 |  |  | YMR170C |  | 1,29168 |
| YOL020W | 448,3277 | 353,2292 |  | YOR222W | 0,44886 | 0,303233 |  | YGR254W |  | 1,228317 |
| YDR035W | 321,6998 |  |  | YPL134C | 0,44886 | 0,303233 |  | YHR174W |  | 1,228317 |
| YHR137W | 318,4523 | 46,12524 |  | **YAL038W (CDC19)** | **0,39342** |  |  | YKL152C |  | 1,228317 |
| YBR117C | 303,4223 | 2,944678 |  | YOR347C | 0,39342 |  |  | YIR031C |  | 1,066515 |
| **YPR074C (TKL1)** | **303,4223** | **2,944678** |  | YOL126C | 0,25278 | 1,357649 |  | YKL060C |  | 1,031041 |
| YJR148W | 250,275 | 200,22 |  | YBR252W | 0 | 15,60595 |  | YLR377C |  | 1,031041 |
| YOR311C | 167,5 |  |  | YDL174C | 0 |  |  | YDR050C |  | 1,029272 |
| YER019W | 143,5891 | 0 |  | YDR272W | 0 |  |  | YER065C |  | 1,028886 |
| YJL121C | 136,8566 | 5,113079 |  | YDR300C | 0 | 1000 |  | YCR012W |  | 1,009732 |
| YPR113W | 122,016 | 363,9394 |  | YEL071W | 0 |  |  | YGR192C |  | 1,009732 |
| YBR029C | 122,0159 |  |  | YKL029C | 0 | 45,94508 |  | YJL052W |  | 1,009732 |
| YDR367W | 109,9168 | 250,25 |  | YML004C | 0 |  |  | YJR009C |  | 1,009732 |
| YKL004W | 109,9168 | 250,25 |  | YOR323C | 0 | 1000 |  | YDR226W |  | 0,546538 |
| YDR072C | 91,72753 | 0 |  | YDL078C |  | 1000 |  | YER091C |  | 0,543448 |
| YDR454C | 61,21212 | 30,96364 |  | YER170W |  | 1000 |  | YGL125W |  | 0,543448 |
| YDR007W | 56,78997 | 42,47148 |  | YGL080W |  | 1000 |  | YPL023C |  | 0,543448 |
| YDR354W | 56,78997 | 42,47148 |  | YGL205W |  | 1000 |  | YDR502C |  | 0,526292 |
| YER090W | 56,78997 | 42,47148 |  | YGR243W |  | 1000 |  | YER043C |  | 0,526292 |
| YGL026C | 56,78997 | 42,47148 |  | YHR002W |  | 1000 |  | YJR105W |  | 0,526292 |
| YKL211C | 56,78997 | 42,47148 |  | YHR162W |  | 1000 |  | YLR180W |  | 0,526292 |
| YGR209C | 45,93409 | 222,1165 |  | YIL160C |  | 1000 |  | YBL039C |  | 0,512229 |
| YDR127W | 7,356177 | 5,589245 |  | YKL188C |  | 1000 |  | YJR103W |  | 0,512229 |
| YGL148W | 7,356177 | 5,589245 |  | YKR009C |  | 1000 |  | YDL022W |  | 0,504723 |
| YBR291C | 6,672427 |  |  | YKR080W |  | 1000 |  | YOL059W |  | 0,504723 |
| YMR241W | 6,672427 | 0 |  | YLR056W |  | 1000 |  | YLR153C |  | 0,488052 |
| YBL068W | 6,473537 | 4,935219 |  | YLR109W |  | 1000 |  | YJL153C |  | 0,458956 |
| YER099C | 6,473537 | 4,935219 |  | YLR284C |  | 1000 |  | YPL087W |  | 0,420455 |
| YHL011C | 6,473537 | 4,935219 |  | YLR348C |  | 1000 |  | YDR287W |  | 0,410628 |
| YKL181W | 6,473537 | 4,935219 |  | YMR015C |  | 1000 |  | YHR046C |  | 0,410628 |
| YOL061W | 6,473537 | 4,935219 |  | YMR272C |  | 1000 |  | YPL061W |  | 0,364362 |
| YER081W | 3,494777 | 5,449761 |  | YNL009W |  | 1000 |  | YER069W |  | 0,27162 |
| YGR208W | 3,494777 | 5,449761 |  | YNL202W |  | 1000 |  | YMR062C |  | 0,27162 |
| YIL074C | 3,494777 | 5,449761 |  | YOR180C |  | 1000 |  | YOL140W |  | 0,27162 |
| YOR184W | 3,494777 | 5,449761 |  | YPL147W |  | 1000 |  | YOR130C |  | 0,27162 |
| YPR021C | 3,10273 | 0,138041 |  | YBL015W |  | 636,3939 |  | YFL030W |  | 0,059875 |
| YPR035W | 2,765313 | 2,189179 |  | YNL117W |  | 500,5333 |  | YNL037C |  | 0,030022 |
| YOR095C | 2,221708 | 5,143796 |  | YDR297W |  | 500,5 |  | YOR136W |  | 0,030022 |
| YDR384C | 1,202505 | 1,032684 |  | YMR165C |  | 500,5 |  | YAL044C |  | 0 |
| YGR121C | 1,202505 | 1,032684 |  | YMR208W |  | 500,2608 |  | YAR035W |  | 0 |
| YNL142W | 1,202505 | 1,032684 |  | YAL054C |  | 500,244 |  | YBR036C |  | 0 |
| YPR138C | 1,202505 | 1,032684 |  | YLR027C |  | 500,221 |  | YBR084W |  | 0 |
| YCR024CA | 1,174844 |  |  | YLR304C |  | 500,022 |  | YBR161W |  | 0 |
| YEL017CA | 1,174844 |  |  | YGR204W |  | 500,0102 |  | YDR019C |  | 0 |
| YGL008C | 1,174844 |  |  | YJR139C |  | 500 |  | YDR148C |  | 0 |
| YPL036W | 1,174844 |  |  | YML126C |  | 500 |  | YER024W |  | 0 |
| YBR196C | 1,138695 |  |  | YNL104C |  | 500 |  | YFL018C |  | 0 |
| YLL052C | 1,108704 | 1,100811 |  | YPL028W |  | 500 |  | YHR144C |  | 0 |
| YPR192W | 1,108704 | 1,100811 |  | YBR183W |  | 364,2727 |  | YIL125W |  | 0 |
| **YLR354C (TAL1)** | **1,071089** |  |  | YLR043C |  | 334,3861 |  | YLR089C |  | 0 |
| YDL085W | 1,070445 | 1,17891 |  | YMR246W |  | 333,6667 |  | YLR174W |  | 0 |
| YMR145C | 1,070445 | 1,17891 |  | YOR317W |  | 333,6667 |  | YML042W |  | 0 |
| YGR248W | 1,064769 |  |  | YGR170W |  | 318,8636 |  | YMR189W |  | 0 |
| YGR256W | 1,064769 |  |  | YOR245C |  | 309,5818 |  | YNL169C |  | 0 |
| YHR163W | 1,064769 |  |  | YCR048W |  | 258,0765 |  | YOR100C |  | 0 |
| YHR183W | 1,064769 |  |  | YNR019W |  | 258,0765 |  | YOR108W |  | 0 |
| YNL241C | 1,064769 |  |  | YMR202W |  | 38,00156 |  | YPL057C |  | 0 |

**Table S5**. FBA results for all pathways in metabolism, including the number of gene targets predicted in each pathway, the total size of each pathway, the fraction of genes in each pathway that are gene targets, and the significance of that representation in each pathway compared to the rest of metabolism (“Whole metabolism”), indicated by a P-value computed with a Fisher's exact test. General pathways such as “carbon metabolism” and “biosynthesis of amino acids” were filtered out of the analysis.

| **KEGG pathway** | **Gene count** | **Pathway size** | **Coverage (%)** | **P-value** |
| --- | --- | --- | --- | --- |
| Pyruvate metabolism | 29 | 47 | 61,70 | 5.64e-10 |
| Pentose phosphate pathway | 18 | 24 | 75,00 | 1.44e-08 |
| Peroxisome | 15 | 21 | 71,43 | 7.54e-07 |
| Valine, leucine and isoleucine degradation | 12 | 19 | 63,16 | 7.19e-05 |
| Glyoxylate and dicarboxylate metabolism | 16 | 30 | 53,33 | 7.73e-05 |
| Glycolysis | 24 | 54 | 44,44 | 8.85e-05 |
| Gluconeogenesis | 24 | 54 | 44,44 | 8.85e-05 |
| One carbon pool by folate | 12 | 20 | 60,00 | 1.47e-04 |
| Aminoacyl-tRNA biosynthesis | 0 | 37 | 0,00 | 2.56e-04 |
| Citrate cycle (TCA cycle) | 16 | 33 | 48,48 | 3.35e-04 |
| Phenylalanine, tyrosine and tryptophan biosynthesis | 11 | 19 | 57,89 | 4.39e-04 |
| Glycine, serine and threonine metabolism | 15 | 33 | 45,45 | 1.57e-03 |
| Lysine degradation | 8 | 13 | 61,54 | 1.67e-03 |
| Starch and sucrose metabolism | 1 | 35 | 2,86 | 4.79e-03 |
| Oxidative phosphorylation | 6 | 70 | 8,57 | 5.71e-03 |
| Nicotinate and nicotinamide metabolism | 0 | 23 | 0,00 | 7.45e-03 |
| Sulfur relay system | 3 | 3 | 100,00 | 9.31e-03 |
| 2-Oxocarboxylic acid metabolism | 14 | 35 | 40,00 | 9.76e-03 |
| Meiosis - yeast | 0 | 21 | 0,00 | 1.20e-02 |
| Galactose metabolism | 0 | 19 | 0,00 | 1.94e-02 |
| Purine metabolism | 19 | 58 | 32,76 | 3.05e-02 |
| Phagosome | 0 | 17 | 0,00 | 3.14e-02 |
| Propanoate metabolism | 9 | 22 | 40,91 | 3.17e-02 |
| Carbapenem biosynthesis | 2 | 2 | 100,00 | 4.44e-02 |
| ABC transporters | 2 | 2 | 100,00 | 4.44e-02 |
| Synthesis and degradation of ketone bodies | 2 | 2 | 100,00 | 4.44e-02 |
| Pyrimidine metabolism | 12 | 33 | 36,36 | 4.71e-02 |
| Whole metabolism | 192 | 909 | 21,12 | - |

**Table S6.** Related to Figure 1. The 30 selected native yeast promoters, and their position in the combinatorial cluster.

| **Number** | **Name** | **Systematic name** | **Position in cluster** |
| --- | --- | --- | --- |
| 01 | pPCK1 | YKR097W | 01 |
| 02 | pTPI1 | YDR050C | 01 |
| 03 | pICL1 | YER065C | 01 |
| 04 | pRNR2 | YJL026W | 01 |
| 05 | pACT1 | YFL039C | 01 |
| 06 | pTDH3 | YGR192C | 01 |
| 07 | pTAL1 | YLR354C | 02 |
| 08 | pENO2 | YHR174W | 02 |
| 09 | pACS1 | YAL054C | 02 |
| 10 | pREV1 | YOR346W | 02 |
| 11 | pCHO1 | YER026C | 02 |
| 12 | pCYC1 | YJR048W | 02 |
| 13 | pTKL1 | YPR074C | 03 |
| 14 | pPGK1 | YCR012W | 03 |
| 15 | pMLS1 | YNL117W | 03 |
| 16 | pBUD6 | YLR319C | 03 |
| 17 | pURE2 | YNL229C | 03 |
| 18 | pCCW12 | YLR110C | 03 |
| 19 | pCDC19 | YAL038W | 04 |
| 20 | pURA1 | YKL216W | 04 |
| 21 | pCRC1 | YOR100C | 04 |
| 22 | pCDC14 | YFR028C | 04 |
| 23 | pTEF2 | YBR118W | 04 |
| 24 | pFBA1 | YKL060C | 04 |
| 25 | pPFK1 | YGR240C | 05 |
| 26 | pTDH2 | YJR009C | 05 |
| 27 | pIDP2 | YLR174W | 05 |
| 28 | pTPK2 | YPL203W | 05 |
| 29 | pRPL15B | YMR121C | 05 |
| 30 | pTEF1 | YPR080W | 05 |

**Table S7.** Related to Figure 1D and 3D. Promoter combinations of library control strains. The numbers in each row refer to promoter numbers as shown in Table S5. Design no. 1 contains the promoters that are native to the genes at the five positions.

| **Design** | **Position 1** | **Position 2** | **Position 3** | **Position 4** | **Position 5** |
| --- | --- | --- | --- | --- | --- |
| 1 | 1 | 7 | 13 | 19 | 25 |
| 2 | 6 | 12 | 18 | 23 | 28 |
| 3 | 4 | 11 | 17 | 24 | 30 |
| 4 | 2 | 8 | 14 | 23 | 28 |
| 5 | 3 | 9 | 15 | 20 | 26 |

**Table S8.** Related to Figure 1 and 4C. Top-30 promoter combinations as recommended by ART. Size of color bars indicate promoter expression strength (see Figure 1), and column “dgfp/dt” shows predicted GFP synthesis rate.

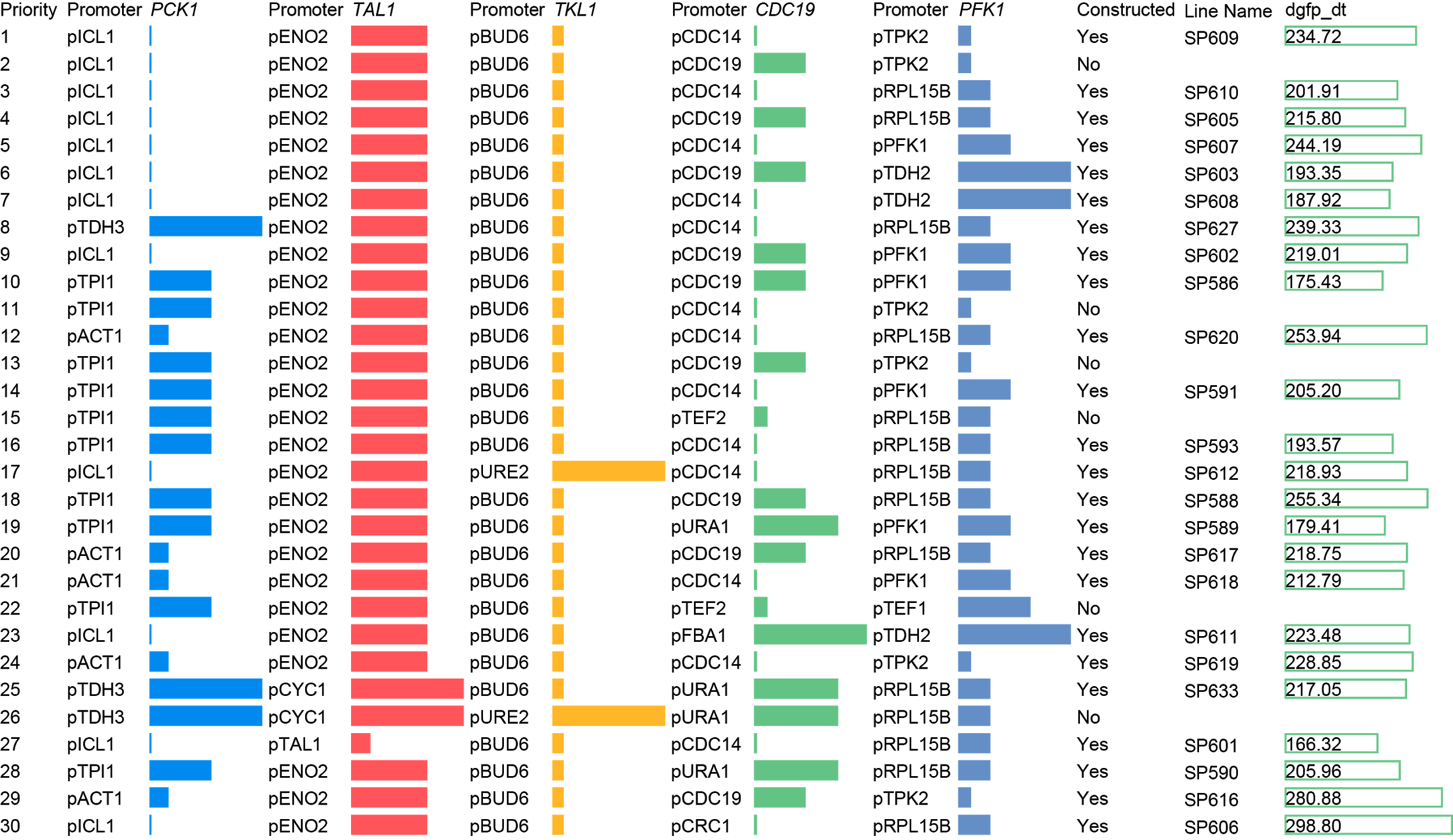

**Table S9.** Related to Figure 1 and 4C. Top-30 promoter combinations as recommended by TeselaGen EVOLVE. Size of color bars indicate promoter expression strength (see Figure 1), and column “dgfp/dt” shows predicted GFP synthesis rate.

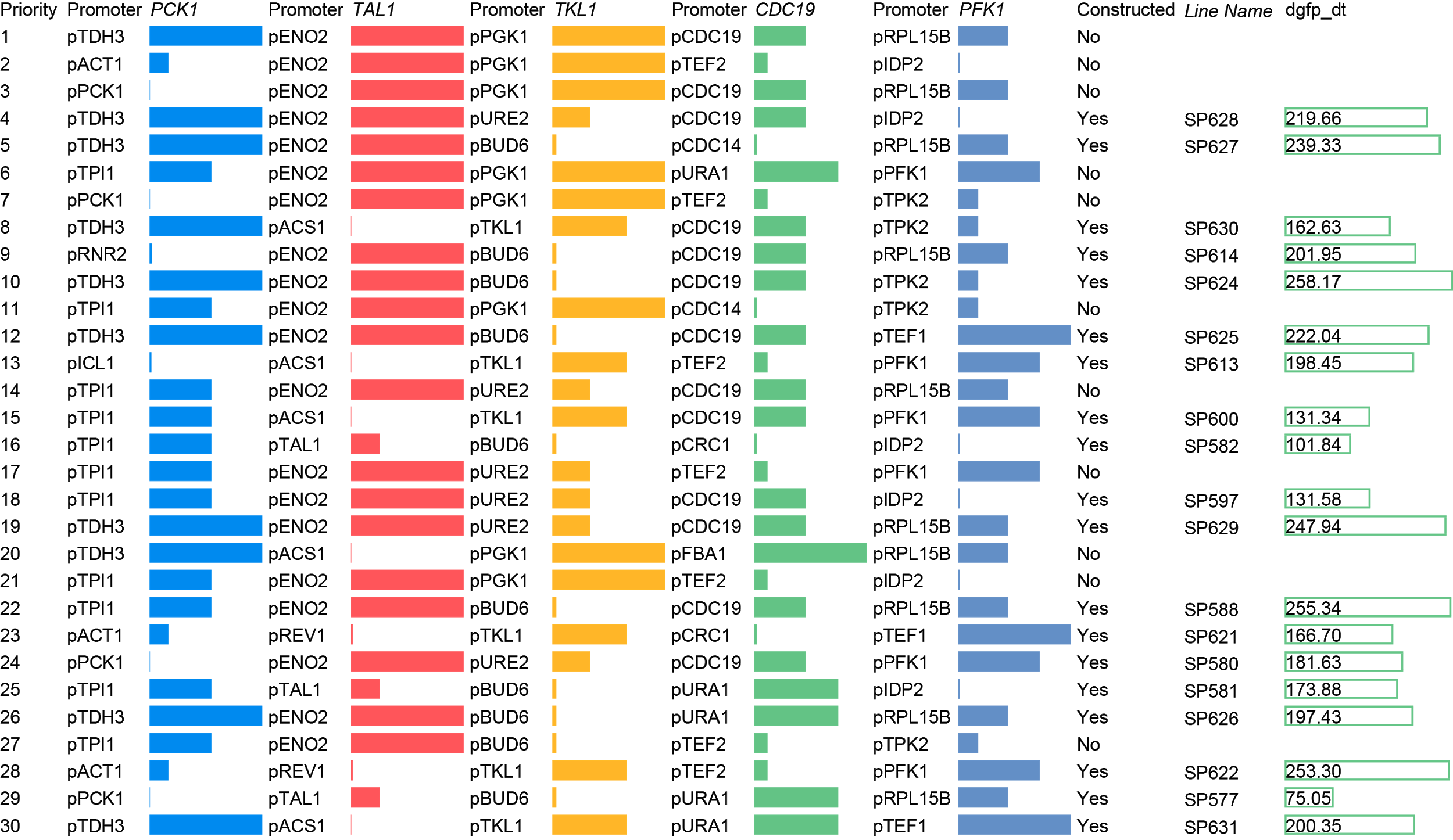
